# Specification of axial identity by Hoxa2 distinguishes between a phenotypic and molecular ground state in mouse cranial neural crest cells

**DOI:** 10.1101/2021.02.09.430457

**Authors:** Irina Pushel, Paul A Trainor, Robb Krumlauf

**Affiliations:** Stowers Institute for Medical Research, Kansas City, MO 64110, USA; Department of Anatomy and Cell Biology, University of Kansas Medical Center, Kansas City, KS 66160, USA

## Abstract

*Hox* genes play a key role in head formation by specifying the axial identity of neural crest cells (NCCs) migrating into embryonic pharyngeal arches. In the absence of *Hoxa2*, NCC derivatives of the second pharyngeal arch (PA2) undergo a homeotic transformation and duplicate structures formed by first arch (PA1) NCCs. Current models postulate that PA1 represents a NCC ‘ground state’ and loss of *Hoxa2* causes a reversion of PA2 NCCs to the PA1 ‘ground state’. We use bulk and single-cell RNAseq to investigate the molecular mechanisms driving this phenotypic transformation in the mouse. In *Hoxa2^-/-^* mutants, PA2 NCCs generally maintain expression of the PA2 transcriptional signature and fail to strongly upregulate a PA1 transcriptional signature. Our analyses identify putative HOXA2 targets and suggest that subsets of NCCs may respond to HOXA2 activity in distinct manners. This separation of phenotypic and molecular states has significant implications for understanding craniofacial development.

## Introduction

Neural crest cells (NCCs) represent one of the defining traits in the evolution of vertebrates (Sauka-Spengler et al. 2007; Green et al. 2015). The neural crest is a transient population of cells that arises along the anterior-posterior axis of the developing central nervous system (Knecht and Bronner-Fraser 2002). NCCs delaminate from the neural epithelium and migrate into the periphery of vertebrate embryos. These migratory and proliferative cells are multipotent and play a dynamic role in vertebrate development, giving rise to diverse structures including neurons, glia, pigment cells, and craniofacial bone and cartilage (LaBonne and Bronner-Fraser 1998; Le Douarin and Kalcheim 1999; Minoux and Rijli 2010). They are divided into trunk and cranial NCCs: trunk NCCs primarily contribute to the peripheral nervous system, while cranial NCCs also give rise to most of the bone and connective tissue of the head and play a key role in craniofacial morphogenesis.

The establishment of cranial NCCs is thought to have been an important step in evolution of the vertebrate head (Northcutt and Gans 1983; Parker et al. 2016; Square et al. 2017). Mutations in genes important for cranial NCC development have been shown to play a role in human craniofacial defects and disorders, making a deeper understanding of NCC development a relevant area of clinical study (Crane and Trainor 2006; Terrazas et al. 2017; Etchevers et al. 2019). Cranial NCCs are initially specified at the neural plate border region, together with neural precursors (LaBonne and Bronner-Fraser 1999; Le Douarin and Kalcheim 1999; Knecht and Bronner-Fraser 2002). The NCCs then delaminate from the dorsal neural tube and migrate into the frontonasal prominence and pharyngeal arches (PAs), highly conserved transient embryonic structures that form the oral apparatus which is essential for feeding and breathing. Upon arrival at their destination, NCCs differentiate into the derivative structures that shape the vertebrate head (Couly et al. 1993)(Fig. 1A,B). NCCs give rise to unique structures based on their axial level of origin (Kontges and Lumsden 1996). For instance, the NCCs migrating into the first PA (PA1) contribute to the maxillary and mandibular components of the jaw, Meckel’s cartilage, and the incus and malleus of the middle ear, while the NCCs migrating into the second PA (PA2) form the stapes of the middle ear and the hyoid bone in the neck (Le Douarin and Kalcheim 1999; Minoux and Rijli 2010).

**Figure 1.**
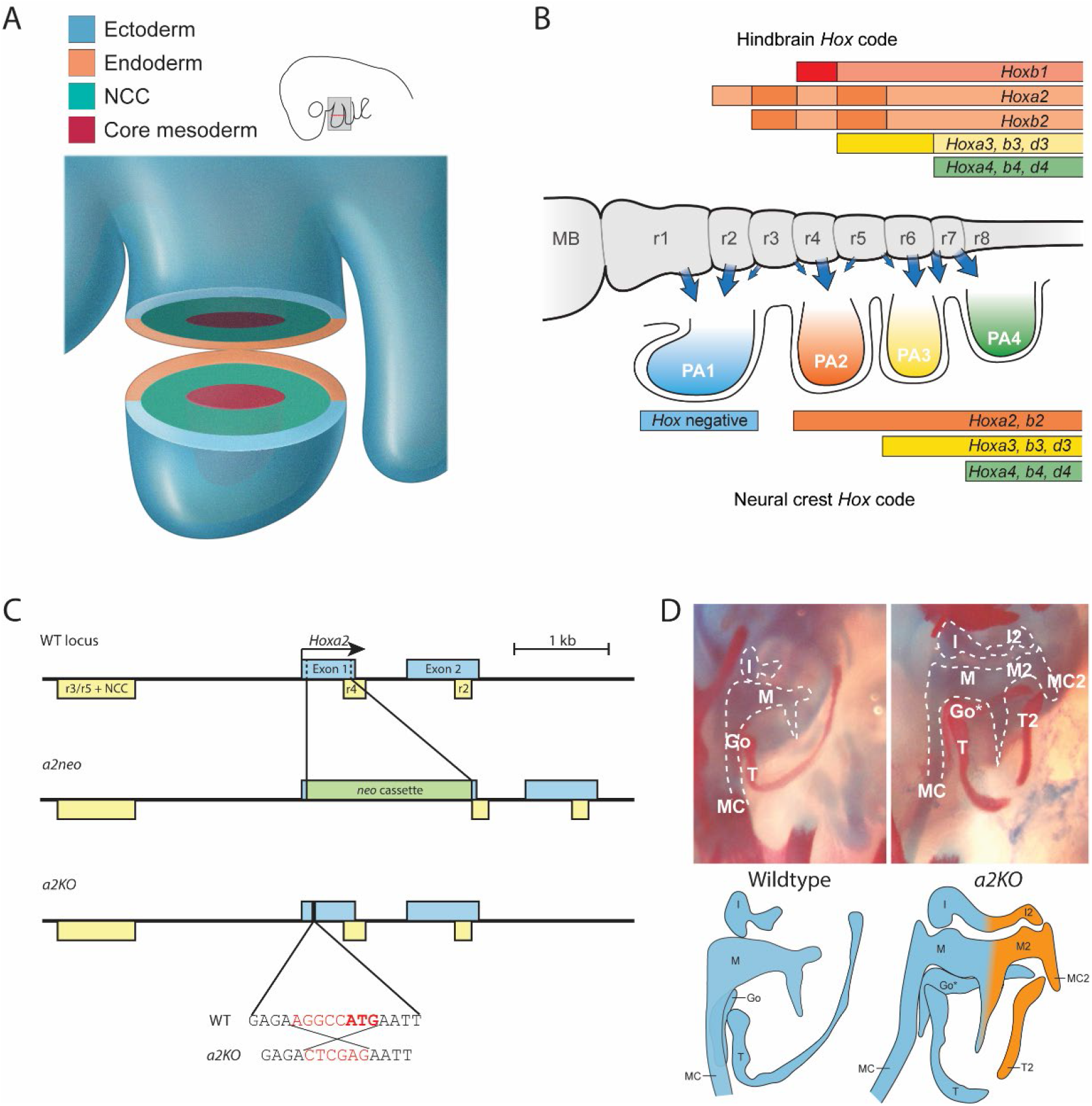
*Hox* genes in the PAs and phenotypic effects of *Hoxa2* knockout. A) NCCs migrate into the PAs surrounding a mesodermal core, contained within the ectoderm and endoderm surrounding the PA structure. Schematic corresponds to ~E9.0-E9.5 mouse embryo. B) *Hox* genes are expressed in a nested pattern that affects NCC fate prior to delamination and migration (hindbrain code, top) and a distinct pattern once the cells have populated the PAs (neural crest code, bottom). Figure adapted from (Parker et al. 2018). C) A comparison of the WT *Hoxa2* locus to the *a2neo* and *a2KO* alleles. Exons depicted in blue, enhancers in yellow. Bottom depicts 7-bp change (highlighted in red) between WT and *a2KO* alleles. Start codon (ATG) bold. D) Comparison of middle ear skeletal staining at E18.5 between embryos that are WT and homozygous for the *a2KO* allele. Cartilagenous structures outlined with white dashed line for clarity. Schematics show PA1-derived structures in blue and PA2-derived structures in orange. M=malleus, I=incus, MC=Meckel’s cartilage, T=Tympanic bone, Go=gonial bone. Duplicated structures designated with a 2 (M2, I2, MC2, T2); Go*=transformed gonial bone.

The highly conserved family of HOX transcription factors plays a key role in establishing regional identities in diverse tissues throughout animal development (Carroll 1995; Alexander et al. 2009; Mallo et al. 2010). During vertebrate head development, the expression and function of *Hox* genes are tightly coupled with the specification of axial identity in hindbrain segments and cranial NCCs (Hunt et al. 1991; Couly et al. 1998; Trainor and Krumlauf 2000b; Trainor and Krumlauf 2001; Minoux and Rijli 2010; Parker et al. 2018). In the developing mouse embryo, NCCs express *Hox* genes in two general phases during their development: prior to their delamination from the neural tube (the ‘hindbrain *Hox* code’) and after they begin migrating into the PAs (the ‘neural crest *Hox* code’) (Fig 1B). This establishes a colinear nested expression pattern of *Hox* genes in the PAs: PA1 NCCs lack *Hox* expression, PA2 NCCs express *Hoxa2* and *Hoxb2*, PA3 NCCs express *Hoxa2, Hoxb2, Hoxa3, Hoxb3*, and *Hoxd3*, and so on into more posterior arches (Hunt et al. 1991; Minoux and Rijli 2010; Parker et al. 2018). This generates a combinatorial *Hox* code that underlies the establishment of unique axial identities of each PA (Trainor and Krumlauf 2001).

Whether *Hox* codes are first established in the hindbrain and transferred into the periphery by migrating NCCs or initiated independently in NCCs separate from the hindbrain *Hox* code is not well understood. Regulatory studies have generated evidence for both independent and shared regulation of *Hoxa2* and *Hoxb2* genes in hindbrain segments and cranial NCCs (Maconochie et al. 1997; Maconochie et al. 1999; McEllin et al. 2016; Parker et al. 2019). Heterotopic grafting experiments in zebrafish and mouse embryos have revealed that *Hox* expression in NCCs is not permanently fixed and is influenced by signals from the PA environment (Trainor and Krumlauf 2000a; Trainor and Krumlauf 2000b; Schilling et al. 2001). This suggests there is dynamic regulation of *Hox* expression during NCC formation and migration in the PAs. However, it remains unclear whether the NCC *Hox* code acts in parallel to or directly within the emerging framework for the conserved core gene regulatory network governing NCC development (Gammill and Bronner-Fraser 2002; Simoes-Costa and Bronner 2015; Martik and Bronner 2017; Parker et al. 2018; Martik et al. 2019).

Loss and gain of function experiments of the *Hox* genes in a number of vertebrate species have led to the formulation of a NCC ground state model for axial patterning of the PAs during embryonic development. In this model, NCCs migrating into the PAs share a common default patterning program, or ground state, that is modified by combinations of *Hox* gene expression to produce distinct derivatives at each axial level (Couly et al. 1998; Minoux and Rijli 2010; Vieux-Rochas et al. 2013). Since NCCs in PA1 lack *Hox* expression, they are postulated to represent the ground state. This hypothesis is largely based on phenotypes associated with perturbation of *Hoxa2* expression. In mouse *Hoxa2^-/-^* mutants, PA2-derived skeletal structures display homeotic transformations and duplicate PA1-derived structures (Gendron-Maguire et al. 1993; Rijli et al. 1993). Conversely, ectopic expression of *Hoxa2* in PA1 NCCs results in transformation of PA1-derived skeletal structures to duplicate PA2 derivatives (Kitazawa et al. 2015). This functional role for *Hoxa2* in patterning PA identity has also been observed in other vertebrate species (Grammatopoulos et al. 2000; Pasqualetti et al. 2000; Hunter and Prince 2002). Together, this evidence supports a conserved role for *Hoxa2* as a master regulator or selector gene that modifies the *Hox*-free NCC ground state to impart a PA2 identity on the NCCs colonizing PA2. However, the precise timing of this *Hoxa2*-dependent process remains unknown, as do the cell populations and downstream target genes through which *Hoxa2* acts to regulate regional identity. There is evidence of a continued requirement for *Hoxa2* activity in NCCs during differentiation (Santagati et al. 2005), but an earlier role for its input into programming the cranial NCCs remains unclear.

In addition to the specific perturbation of *Hoxa2* expression, deletion of the entire *HoxA* cluster in the mouse results in PAs 2-4 all producing additional PA1-like structures in lieu of caudal derivatives. This further supports the existence of a *Hox*-free ground state that is transformed in each of the PAs by distinct combinations of *Hox* gene expression (Minoux et al. 2009). It is clear from the phenotypic transformations observed in mutant embryos that there exists a phenotypic NCC ground state represented by the set of NCC derivatives produced in the absence of *Hox* gene expression. Whether this phenotypic ground state is accompanied by a shared molecular ground state – a transcriptional signature – that corresponds to *Hox*-free NCCs and is observed in *Hox* mutant embryos, has not yet been addressed. Where the term ‘ground state’ has been used to date, the explicit distinction between a phenotypic and molecular ground state has not been made (Rijli et al. 1993; Minoux et al. 2009; Amin et al. 2015). This is an important issue to resolve for understanding regulatory mechanisms that pattern craniofacial development, because *Hoxa2* could be working to modify the properties of all NCCs in PA2 or only sub-populations of cells required for specifying PA2 derived structures. Regardless of the process, *Hoxa2* clearly has a fundamental effect in determining PA2 identity.

Recent advances in sequencing technologies enable us to more thoroughly explore the molecular underpinnings of axial specification and distinguish between phenotypic and transcriptional effects at single-cell resolution. Several studies have provided comparative analysis of gene expression in the PAs (Brunskill et al. 2014; Lumb et al. 2017; Minoux et al. 2017). However, these comparisons have not yet been extended to *Hox* mutant embryos, and thus lack the power to determine whether a *Hox*-free mutant PA (other than PA1) reverts not only at the phenotypic level but also at the molecular or transcriptional level to a PA1-like state. Thus, elucidating the molecular mechanism by which *Hoxa2* expression acts to impart a PA2 NCC fate directly informs our understanding of the transcriptional basis of axial identity specification in the developing mouse embryo.

In this study, to deepen our understanding of the role of *Hoxa2* in regulating PA2 identity, we have used bulk and single cell RNAseq in mice, comparing wildtype (WT) and *Hoxa2* mutant embryos, to investigate the precise timing and populations of cells where it exerts its regulatory activity on NCCs. Strikingly, our data do not show a global reversion of the *Hoxa2^-/-^* PA2 transcriptional profile in NCCs to a PA1-like state, as would be expected in the presence of a molecular NCC ground state. Rather, we find evidence that *Hoxa2* exerts its regulatory activity in distinct subpopulations of cells in PA2 as NCCs are undergoing differentiation. Using differential expression analyses from transcriptional profiling of cell populations and single cells, we identified both previously described and novel putative targets of *Hoxa2* involved in PA2 fate specification. Taken together, our data suggest that the phenotypic ground state of mouse cranial NCCs is not matched with an underlying molecular ground state and reveal novel downstream targets of *Hoxa2* involved in axial identity specification. Our findings highlight the value of scRNA-seq in evaluating and refining models surrounding the molecular basis of developmental and evolutionary phenotypes. Moreover, the rich transcriptomic datasets generated in this study are a valuable resource to further characterize the molecular underpinnings of NCC axial identity.

## Results

### Generation of a new mouse *Hoxa2* null allele

In order to investigate the molecular mechanisms by which *Hoxa2* expression drives PA2 fate, we used a strategy based on transcriptional profiling approaches to compare wildtype (WT) and *Hoxa2* mutant mouse embryos. Existing mouse lines have mutations with a *neomycin* cassette inserted into *Hoxa2* (Gendron-Maguire et al. 1993; Rijli et al. 1993), which may alter the expression of neighboring *Ho*x genes, as we know this is a region rich in regulatory elements both within and flanking the gene (Tümpel et al. 2006) (Fig. 1C). These lines also exist on a mixed genetic background and we wanted to minimize noise in genomic comparisons by performing analyses in a consistent background. In addition, we wanted to be able to monitor endogenous *Hoxa2* mRNA expression in cells lacking functional HOXA2 protein. To do this, we generated a novel null allele by using CRISPR-Cas9 gene editing to alter a 7 bp region in the endogenous locus which includes part of HOXA2 start codon, deleting 2 bp and converting it into an *XhoI* site (5’-<u>AGGCC</u>**<u>AT</u>G-**3’ to 5’-<u>CTCGA</u>**G**-3’). We refer to this allele as *a2KO*, which minimizes changes to the locus and preserves all known *cis*-regulatory elements (Fig. 1C).

To validate the successful abrogation of HOXA2 protein function by this mutation, we compared homozygous *a2KO* embryos to a previously characterized mutant containing the *neomycin* cassette (Rijli et al. 1993) (Fig. 1C-D). We refer to this published allele as the *a2neo*. As with pups homozygous for *a2neo*, homozygous *a2KO* pups die within 24 hours of birth and display the previously characterized ‘cauliflower ear’ phenotype. Importantly, *a2KO* embryos also show the same duplication of PA1-derived middle ear elements that is characteristic of the *a2neo* embryos at E18.5 (Fig. 1D). It is interesting to note that *a2KO* embryos do not show the cleft palate defect originally observed in *a2neo* mice (Rijli et al. 1993). Our *a2neo* colony also lacks cleft palate, and a complementation test intercrossing *a2KO* and *a2neo* heterozygous alleles also fails to recapitulate the cleft palate phenotype. These observations may be due to differences in genetic background or an off-target effect of the *neomycin* cassette disrupting expression of other genes in the *HoxA* cluster. Homozygous embryos with *a2KO* or *a2neo* alleles show matching phenotypes with respect to PA2 derived structures and affected tissues, rendering the *a2KO* allele a valid genetic model for studying *Hoxa2* function in NCCs.

### PA1 and PA2 have distinct molecular states early in NCC migration

The ‘ground state’ model considers that *Hoxa2* is a selector gene for PA2 fate, modifying the *Hox*-free ground state observed in PA1 (Fig. 1B). Thus, we wanted to understand when differences appear in the transcriptional profiles of NCCs from PA1 and PA2 during development and when HOXA2 acts to modify the putative ground state to specify PA2 fates. To address these questions, we dissected PA1 and PA2 from wildtype (WT) and homozygous *a2KO* mutant embryos at four timepoints, ranging from early NCC migration (E9.0) to the start of differentiation (E10.5), and examined their transcriptional profiles by bulk RNAseq (bRNAseq). While NCCs represent the majority of cells in the dissected PA tissue, the arches also contain cells from the surface ectodermal and endodermal layers and a core of mesodermal cells (Fig. 1A). We utilized the entire PA tissue for these experiments to avoid inducing changes in cell properties during manipulations or finer level dissections and to ensure we had the full cellular repertoire of components of the PAs.

We first examined when molecular differences between PA1 and PA2 appear during NCC migration and early differentiation by performing differential gene expression analysis (Supplementary Tables S1-5). If early NCCs migrating into the PAs have a common ground state that is progressively modified in PA2 by HOXA2, their initial transcriptional profiles in PA1 and PA2 would be predicted to be quite similar, with differences emerging and becoming more pronounced as HOXA2 exerts its activity during the course of NCC colonization of the PAs. Our data reveal that early in migration at E9.0, PA1 and PA2 already display significantly different transcriptional profiles and this pattern persists through E9.5 (Fig. 2A), arguing against a common molecular ground state early in the developmental process. Then as the embryo continues to develop from E10.0-E10.5, at the onset of NCC differentiation, we observe progressively fewer differences between PA1 and PA2. The majority of the differentially expressed genes are unique to a particular timepoint, although 89 genes are PA-biased throughout development and a fair number of genes are differentially expressed at both E9.0 and E9.5, when we observe the greatest transcriptional differences between PA1 and PA2 (Supplementary Fig. S1A-B).

**Figure 2.**
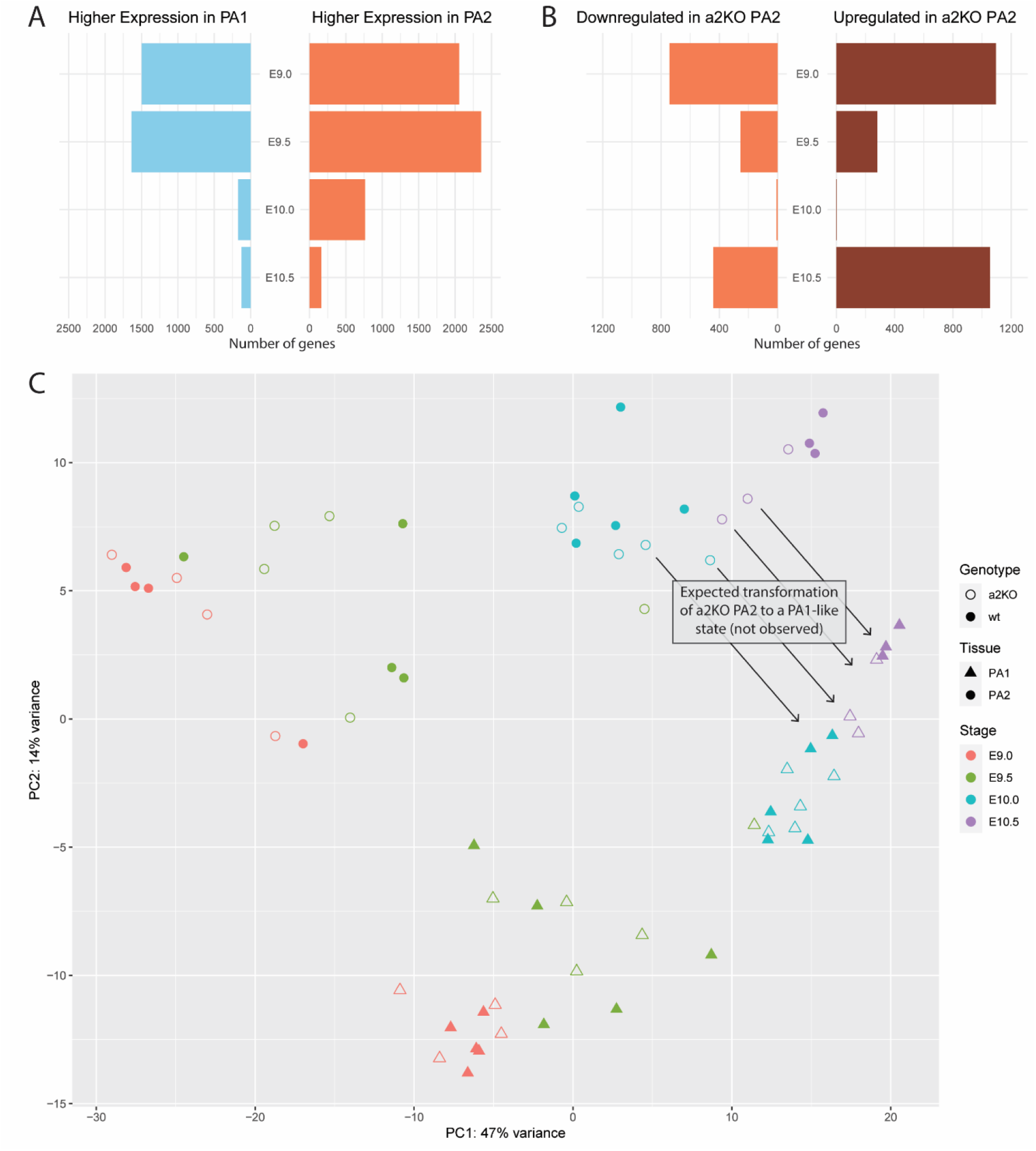
Hoxa2 knockout PA2 does not revert to a PA1-like transcriptional state. A) Number of genes differentially expressed between WT PA1 and WT PA2 at each timepoint E9.0-E10.5. B) Number of genes differentially expressed between WT PA2 and a2KO PA2 at each timepoint E9.0-E10.5. C) Principal component analysis (PCA) plot depicting clustering of WT PA1, WT PA2, a2KO PA1, and a2KO PA2 from E9.0-E10.5. Each point corresponds to a sample from an individual embryo (PA1 and PA2 were both collected from each embryo). Color corresponds to timepoint, shape corresponds to tissue, fill corresponds to genotype. All samples cluster according to PA of origin, with PA2 in upper left of the plot and PA1 in lower right, independent of genotype.

Overall, these data indicate that cranial NCCs emerging from the hindbrain at different axial levels have distinct transcriptional profiles as they begin to migrate into PA1 and PA2, suggesting that axial identities may already be established by early stages of NCC migration. The similarity of transcriptional profiles in both arches at later stages may reflect that NCCs are undergoing differentiation programs that generate a similar repertoire of cell types - osteoblasts, chondrocytes, neurons, glia, and melanocytes in each pharyngeal arch environment. These cells may exist in different proportions and end up in unique orientations relative to one another, ultimately resulting in distinct derivatives in PA1 and PA2, however the cell types forming these structures are similar.

### *Hoxa2* maintains the molecular character of PA2 in different phases of NCC development

To investigate how the loss of HOXA2 affects PA2 development, we compared the transcriptional profiles between PA2 of WT and *a2KO* embryos and identified genes differentially expressed during NCC migration and early differentiation (Fig. 2B and Supplementary Tables S6-10). At the earliest timepoint (E9.0), there are significant transcriptional differences in PA2 between WT and *a2KO* embryos, indicating that *Hoxa2* plays an important early role in shaping PA2 identity. These molecular differences then decrease as NCCs colonize the PAs (E9.5-E10.0), but then pronounced differences once again arise in their expression profiles as they begin to differentiate at E10.5 (Fig. 2B). Many of the genes differentially expressed between WT PA2 and *a2KO* PA2 are specific to a particular timepoint. For example, 65% of genes differentially expressed at E9.0 and 62% of genes differentially expressed at E10.5 are not differentially expressed at any other timepoint (Supplementary Fig. S1C-D). Thus, the loss of HOXA2 results in series of dynamic changes in the transcriptional program of NCCs from PA2. Consistent with the observed *Hoxa2^-/-^* phenotype of increased ossification in PA2, the genes differentially expressed at E10.5 include *Msx2*, which is expressed at higher levels in *a2KO* PA2 than WT PA2, suggesting an increase in osteogenesis. Taken together, these data suggest that *Hoxa2* has multiple roles in regulating transcriptional programs in different phases of NCC development - i.e. migration and differentiation.

### PA2 is not transformed to the molecular character of PA1 in the absence of *Hoxa2* activity

Our bRNAseq analysis indicates that the establishment and maintenance of the transcriptional signature of PA2 depend upon HOXA2 (Fig. 2B). We then examined whether the changes observed in the transcriptional profiles of PA2 represent a general reversion to that of PA1, as predicted by the ground state model. To compare transcriptional signatures over the timecourse between the WT and *a2KO* samples, we performed principal component analysis (PCA) to visualize similarities and differences between their profiles (Fig. 2C). WT PA1 and PA2 begin by clustering separately at E9.0 and E9.5, in accordance with our observation that in these early stages of NCC development, gene expression differs greatly between the two (Fig. 2A, C). The differences are less pronounced at the E10.0 and E10.5 timepoints. If PA2 converts to a PA1-like signature in *a2KO* mutants, in association with the transformation of PA2 derivatives to PA1, at some point in the timecourse we expected to see *a2KO* PA2 begin to cluster with PA1 of WT and *a2KO* embryos. To our surprise, the transcriptome of *a2KO* PA2 samples at all timepoints continues to cluster with the corresponding transcriptome of WT PA2and do not at any point cluster with PA1 samples from WT or *a2KO* embryos (Fig. 2C). The fact that this continues into E10.5, when gene activity has previously been shown to be altered due to the absence of *Hoxa2* (Santagati et al. 2005; Donaldson et al. 2012), suggests that the clear phenotypic transformation of PA2 derivatives in *a2KO* embryos is not accompanied by a general transition in the transcriptional state of PA2 to that of PA1. Hence, the phenotypic ground state is not reflected in an underlying common molecular ground state.

### Single-cell transcriptomics reveals heterogeneity of PA1 and PA2 during NCC differentiation

One possible explanation for the observed lack of transformation at the transcriptional level is heterogeneity in the tissue collected via manual dissection of PAs (Fig. 1A & 3B). While a large fraction of the cells in the PAs at the stages examined are NCCs, cells from other tissues of the PAs (ectoderm, endoderm and mesoderm) may make it difficult to identify NCC-specific effects using whole PAs. The concerns about tissue heterogeneity and the challenges of understanding tissue-specific effects of *Hoxa2* in this context led us to pursue a higher-resolution technique to address these questions at the level of single cells. Because our bRNAseq analysis and previously published data (Santagati et al. 2005; Donaldson et al. 2012; Amin et al. 2015) support a significant contribution of HOXA2 to NCC differentiation, we collected PA1 and PA2 from WT and *a2KO* embryos at E10.5 and performed scRNAseq to compare gene expression at the single-cell level (Fig. 3A). As with our bRNAseq experiments, we used whole PAs to maintain cell health during collection and to avoid biasing the isolation process by ensuring access to a full repertoire of cellular components of the PAs. The presence of non-NCC tissues also allows further interrogation of cell non-autonomous effects. After filtering the data for quality, we performed an integrated analysis in Seurat v3 (Stuart et al. 2019) to concurrently look at all four samples (Supplementary Fig. S2). To facilitate accessibility to these data, we have created and made available an R ShinyApp to visualize gene expression across cells in this dataset, available at https://simrcompbio.shinyapps.io/irp_scrnaseq_2020/.

**Figure 3.**
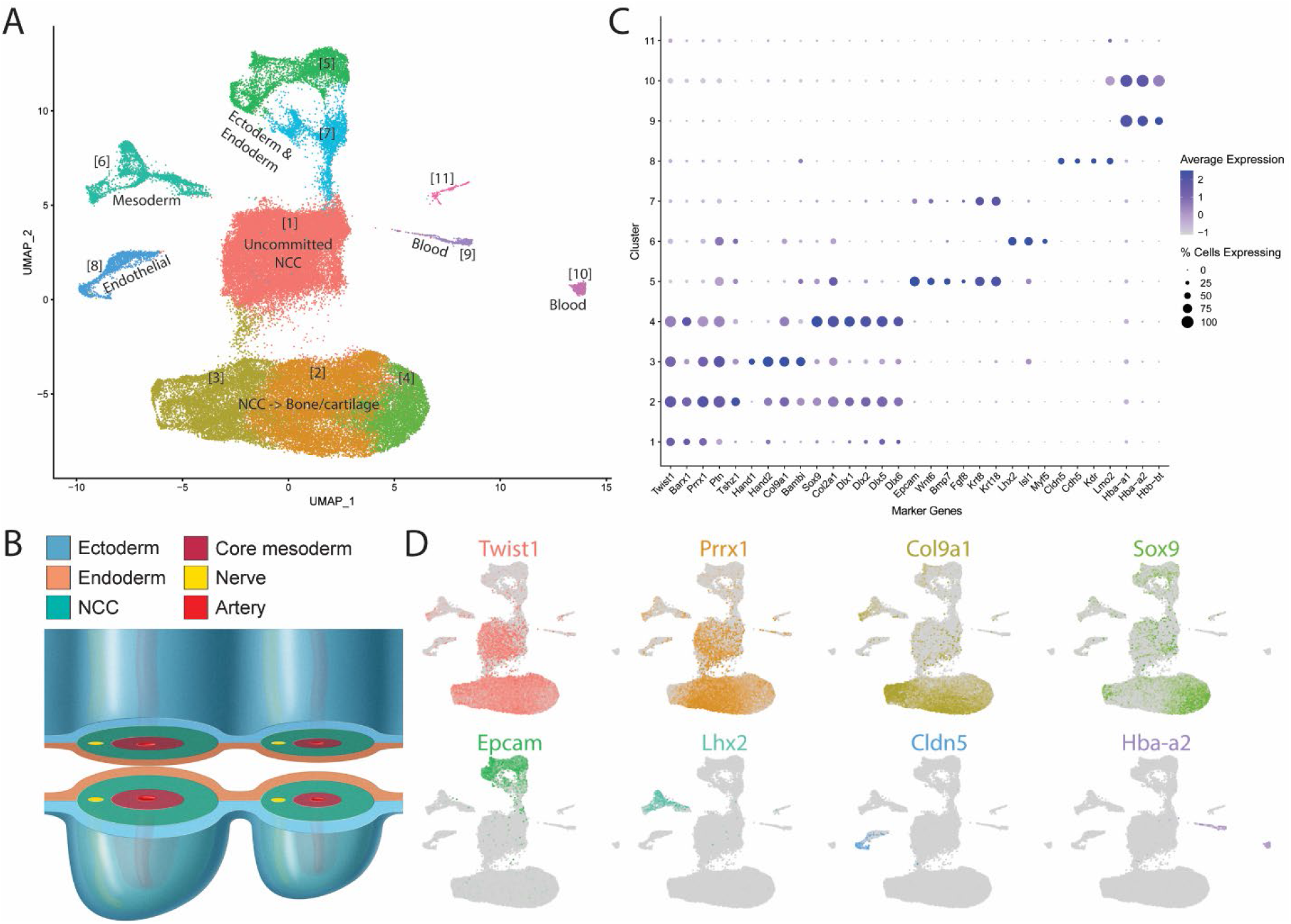
Single-cell RNAseq data demonstrate the heterogeneity of cells in the PAs. A) Uniform manifold approximation and projection (UMAP) depicting a total of 67,674 cells across the four samples were used for downstream analysis after quality control filtering for low sequencing depth and doublets. B) Cross-section representing key PA structures at the onset of NCC differentiation (E10.5). C) Marker gene expression allows the identification of eleven distinct clusters common to all PAs sampled. D) UMAP visualization of marker genes for selected clusters.

Clustering the cells from all four samples results in 67,674 cells arranged in 11 clusters (Fig. 3A). Based on known marker genes characteristic of tissues and cell types, we were able to assign identities to the cells in each cluster (Table 1, Fig. 3C-D, Supplementary Table S11).

**Table 1.** Cluster identity of cells from E10.5 WT PA1, WT PA2, a2KO PA1, and a2KO PA2.

| Cluster | Identity | Marker genes |
| --- | --- | --- |
| 1 | Uncommitted NCC | *Twist1*, *Barx1*, *Crabp1*, *Dlx6* |
| 2 | NCC -> bone/cartilage | *Twist1*, *Prrx1*, *Col9a1* |
| 3 | NCC -> bone/cartilage | *Hand1*, *Hand2*, *Col9a1* |
| 4 | NCC -> bone/cartilage | *Sox9*, *Col2a1*, *Dlx1*, *Dlx2*, *Barx1* |
| 5 | Endoderm & ectoderm | *Epcam*, *Wnt6*, *Fgf8*, *Bmp7* |
| 6 | Mesoderm | *Myf5*, *MyoD* |
| 7 | Endoderm & ectoderm | *Epcam*, *Wnt6*, *Fgf8*, *Bmp7* |
| 8 | Endothelial | *Cldn5*, *Kdr*, *Lmo2* |
| 9 | Blood | *Hbb-bs*, *Hba-a1*, *Hbb-bt* |
| 10 | Blood | *Hba-bh1*, *Hba-a1*, *Hbb-a2* |
| 11 | Unknown | *Pf4*, *Fcer1g*, *Tyrobp*, *C1qb*, *Lyz2* |

At E10.5, the majority of the cells (80.2%), represented by clusters 1-4, are of neural crest origin and many of them show gene expression profiles characteristic of cells that have begun to differentiate into NCC derivatives. Clusters 2-4 correspond to NCCs differentiating into bone and cartilage, characterized by the expression of established marker genes such as *Sox9, Col2a1, Col9a1, Prrx1, Hand1*, and *Hand2*. We also observe a population of NCCs (cluster 1) that express known NCC markers, including *Twist1* and *Barx1*, but do not express genes indicative of a commitment to a particular fate. These may represent a later-migrating population of NCCs that have yet to initiate programs of differentiation at the time of tissue collection.

In addition to NCCs, we also observe populations of ectodermal and endodermal cells (clusters 5 & 7), as well as pharyngeal mesoderm (cluster 6) and endothelium (cluster 8). Two clusters of cells (clusters 9 & 10) are largely defined by their expression of hemoglobin genes, suggesting that they are blood cells. There are several possible sources of blood cells in this experiment that likely explain the presence of these two clusters - one source being contamination picked up from the dish during PA isolation and the other corresponding to blood cells developing in the PAs. Finally, there is a small cluster (11) of cells that do not correspond to a known cell type. In summary, these transcriptional profiles of single cells within the PAs allow us to identify populations that reflect specific tissues and cell types known to reside within PA1 and PA2 of WT and *a2KO* embryos, including NCCs undergoing differentiation.

### NCC clustering reflects spatial, temporal and cell type differences

The scRNAseq dataset enables us to identify individual subsets of NCCs as they commit to specific fates and begin to form derivative structures. The four clusters of NCCs and NCC derivatives (clusters 1-4) display underlying transcriptional differences that reflect spatial, temporal and cell type differences between and within PA1 and PA2. For example, looking at the distribution of proximal-distal markers, we see a clear gradient through the bone and cartilage derivatives (clusters 2-4). Cluster 3 contains distal NCCs marked by expression of *Hand1* and *Hand2*, while cluster 4 corresponds to proximal NCCs marked by expression of *Barx1* and *Pou3f3* (Fig. 4A). This data suggests a clear proximal-distal (right-left) order is reflected in the clustering of NCC-derived bone/cartilage. In cluster 1, the less committed NCCs, we see lower levels and greater heterogeneity in the expression of these proximal-distal markers, supporting the idea that this cluster corresponds to NCCs found throughout the arch that have not yet adopted a particular fate. Examining rostral-caudal markers in clusters 2-4, we see that cells in the lower left are marked to a greater extent by *Gsc*, indicating a caudal bias, while cells in the upper right are marked by *Lhx8*, indicative of rostral cells (Fig. 4B). Cluster 1 again shows lower levels and more heterogeneous expression of rostral-caudal markers, consistent with the idea that these cells cluster according to their identity as less committed NCCs rather than based on a spatial location within the arches. This illustrates that clustering trends that generate clusters 2-4 have captured positional information and spatial arrangements of NCCs in the PAs.

**Figure 4.**
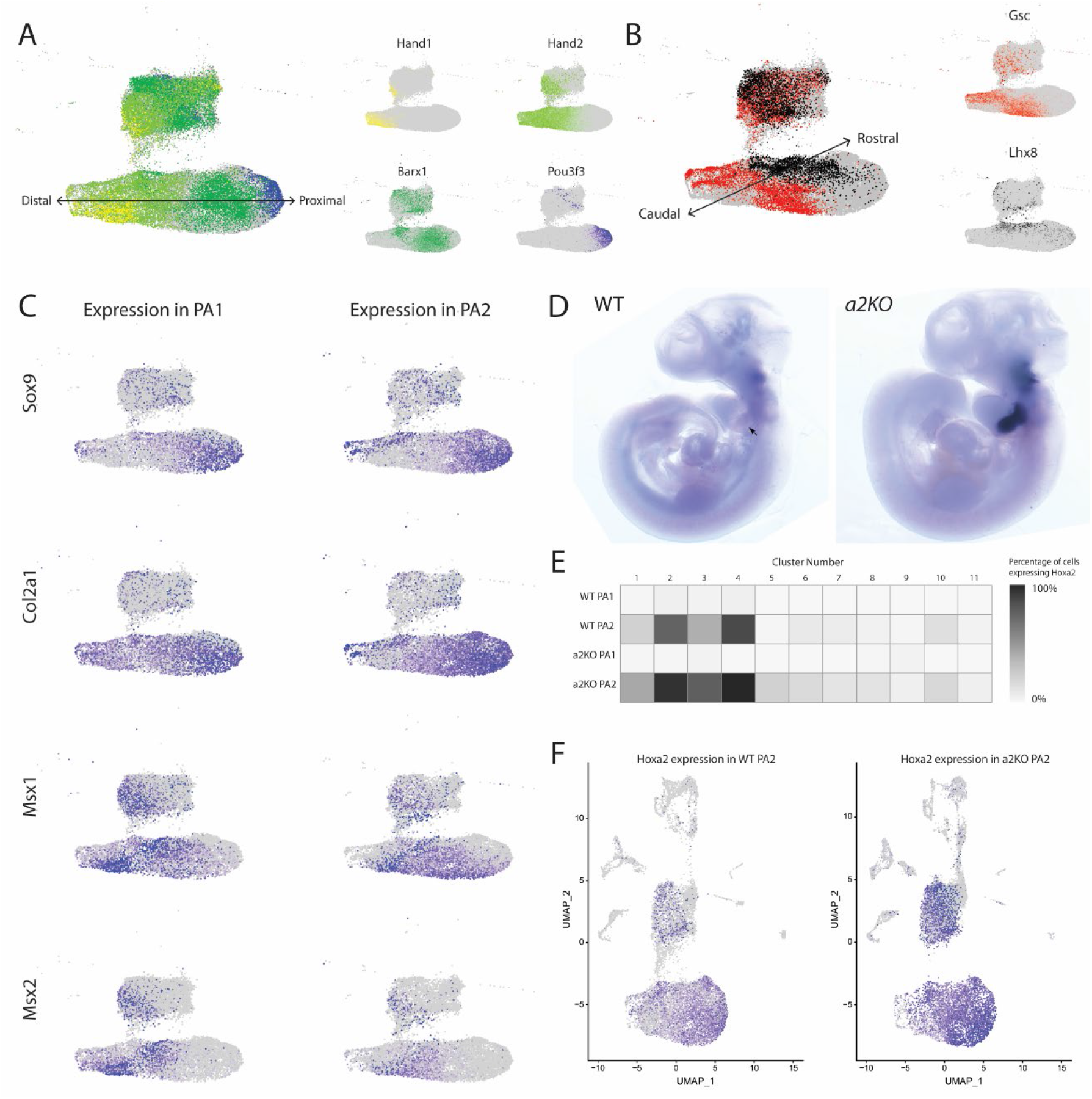
Spatial markers and *Hoxa2* expression delineate subsets of NCC derivatives. A) A left-right distal-proximal axis emerges in NCC-derived bone/cartilage of the PAs. *Hand1* and *Hand2* expression mark distal cells, while *Barx1* and *Pou3f3* expression mark proximal cells. B) An upper right-lower left rostral-caudal axis emerges in bone/cartilage NCC derivatives, marked by *Lhx8* (rostral) and *Gsc* (caudal) expression. C) Markers of chondrogenesis, *Sox9* and *Col2a1*, are segregated in their expression from markers of osteogenesis, *Msx1* and *Msx2*. D) *Hoxa2* expression in WT and *a2KO* mouse embryos at E10.5. D) *Hoxa2* expression in WT PA2 (left) and *a2KO* PA2 (right). *Hoxa2* transcript is still observed in a2KO PA2 due to the nature of the *a2KO* allele, in which the gene is still transcribed but the transcript cannot be translated. E) Percentage of cells in each cluster expressing *Hoxa2* transcript. F) Visualization of *Hoxa2* expression in WT PA2 (left) and *a2KO* PA2 (right) in scRNAseq data.

With respect to cell types, markers of chondrogenesis and osteogenesis also display a fair amount of segregation (Fig. 4C and Supplementary Fig. S3). *Sox9* and *Col2a1*, hallmarks of chondrogenesis, show the highest levels of expression in cluster 4, with limited expression at the distal-most tip of cluster 3. We observe similar levels of expression between PA1 and PA2, with a bias toward the proximal domain of the arches, consistent with previous descriptions of their expression domains. In contrast, *Msx1* and *Msx2*, regulators of osteogenesis, are largely expressed in cluster 2. Expression of these genes is higher in PA1 than PA2, as expected considering greater intramembranous ossification in PA1 (Minoux and Rijli 2010). Moreover, *Msx1* and *Msx2* expression is biased in both arches toward the distal end of the clusters, consistent with their endogenous patterns of expression. This segregation and lack of overlap between differentiation markers supports the observation that chondrogenesis and osteogenesis are occurring in distinct groups of cells. In cluster 1, the expression of all of these markers is lower and less segregated, consistent with the idea that these cells have not yet committed to a bone/cartilage fate. Taken together, these transcriptional profiles reveal that NCC-derived bone and cartilage cells, undergoing differentiation at E10.5, show spatial and cell differentiation biases in their clustering that capture diverse endogenous features of NCCs in the PAs.

### *Hoxa2* expression is enriched in NCCs and NCC-derived clusters at E10.5

To probe the role of *Hoxa2* in specifying axial identity, we next wanted to examine its expression within the scRNAseq dataset. We designed the *a2KO* mutation so that the homozygous mutant mice produce a *Hoxa2* transcript with a small change (7 nt) spanning the initiation codon, which prevents translation. This enables us to monitor endogenous *Hoxa2* transcripts in embryos of both WT and *a2KO* embryos. To validate this ability, we performed whole mount in situ hybridization in E10.5 WT and *a2KO* embryos and found that *Hoxa2* is expressed in identical domains in the hindbrain and PAs (Fig. 4D). Intriguingly, we observe higher levels of *Hoxa2* transcripts in PA2 of the *a2KO* mutants compared to WT embryos, which is consistent with levels observed in our bulk and single cell RNAseq data (Fig. 4F). This domain-specific elevation of *Hoxa2* transcript levels may arise through the input of cross-regulatory feedback circuits known to modulate *Hox* expression in the hindbrain and NCCs (Parker et al. 2019; Parker and Krumlauf 2020). The ability to monitor endogenous *Hoxa2* transcripts in *a2KO* mutants is important because the expression of *Hoxa2* serves as a lineage tracer in this system and indicates that these cells survive even in the absence of HOXA2. Moreover, directly comparing the properties of cells expressing *Hoxa2* in WT and *a2KO* embryos enables us to identify transcriptional differences associated with *Hoxa2*-dependent activity.

In PA1 of WT and *a2KO* embryos, which are expected to be *Hox*-negative, fewer than 5% of the cells express *Hoxa2* and they are not enriched in any single cluster (Fig. 4E and Supplementary Fig. S4). The low fraction of *Hoxa2*-expressing cells observed could be due to contamination during the cell collection process or a small population of *Hoxa2*-expressing cells in the tissue that have not been previously described. In both WT and *a2KO* PA2, we see the highest levels of *Hoxa2* transcript in NCCs and NCC-derived clusters (1-4), with very few cells expressing *Hoxa2* in surface ectoderm, endoderm, mesoderm or endothelia (Fig. 4F). Expression of *Hoxa2* is spread throughout clusters 2-4, with higher levels of expression in clusters 2 and 4 than clusters 1 and 3. The differences in expression levels in NCC clusters may be related to the emerging identities and fates of these cells and the temporally dynamic roles of HOXA2 on the establishment of these identities.

### Comparing transcriptional signatures of PA1 and PA2

We sought to leverage the richness of our bRNAseq and scRNAseq datasets to understand key characteristics that define the transcriptional states of PA1 and PA2, as well as how these states are modified in the absence of *Hoxa2*. Returning to the differential expression analysis of our bRNAseq datasets, we focused on early differentiation at E10.5 and looked at genes differentially expressed between WT PA1 and WT PA2 (Supplementary Table S12). We refer to the 126 genes more highly expressed in PA1 as the ‘PA1 transcriptional signature’ and the 164 genes more highly expressed in PA2 as the ‘PA2 transcriptional signature’ (Fig. 5A-B).

**Figure 5.**
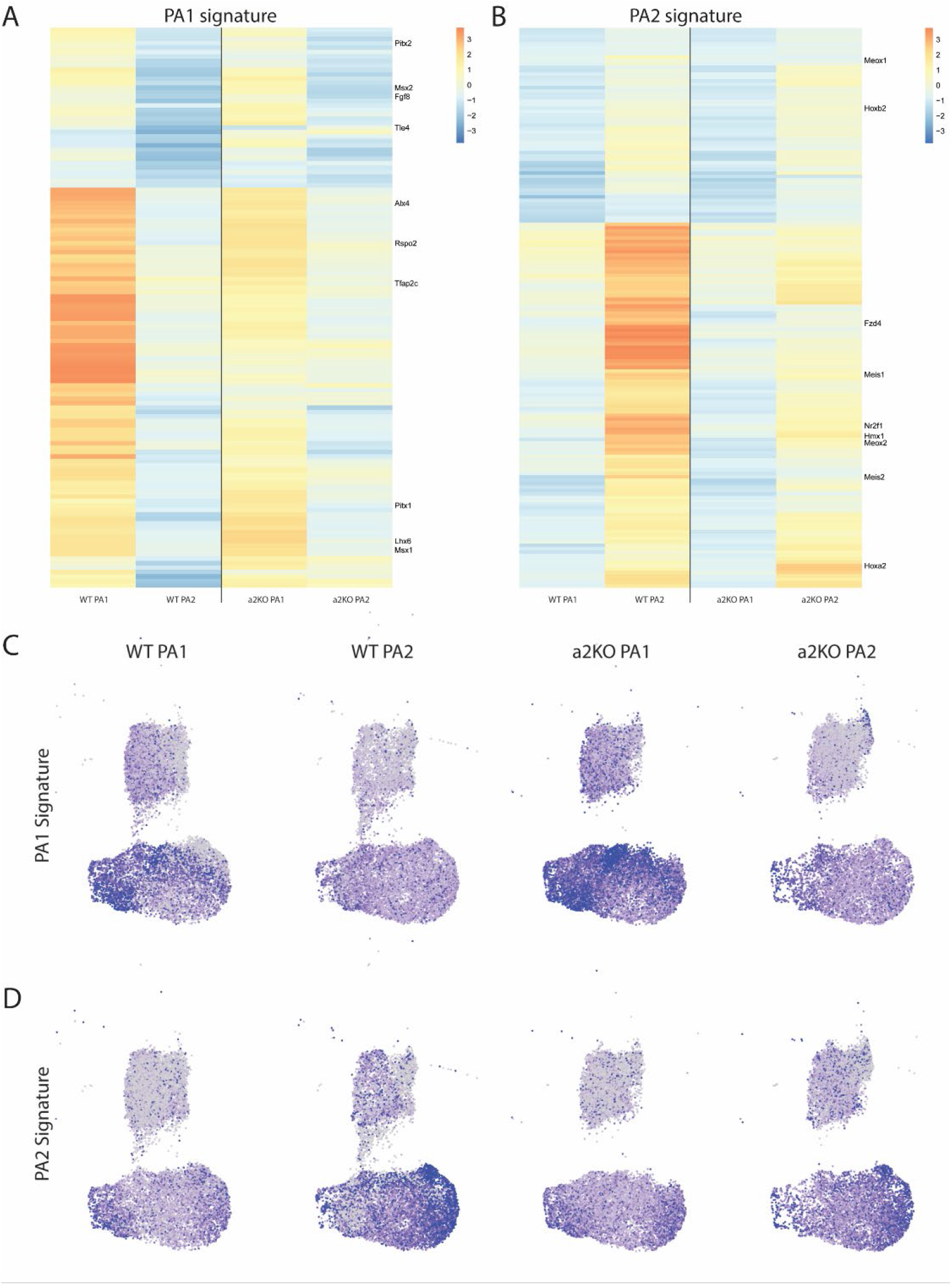
Transcriptional signatures of PA1 and PA2 at the onset of NCC differentiation. Differential expression analysis from bRNAseq between WT PA1 and WT PA2 at E10.5 generated a 126-gene PA1 transcriptional signature and 164-gene PA2 transcriptional signature. Visualization of these signatures across bRNAseq (A & B) and scRNAseq (C & D) E10.5 datasets across WT PA1, WT PA2, *a2KO* PA1, and *a2KO* PA2 reveals trends in transcriptional identity. bRNAseq data (A & B) shown as Z-scores of expression across all conditions and timepoints for each gene. scRNAseq data (C & D) shown as normalized expression of a MetaFeature for each PA-specific signature across NCCs in each condition.

We then compared the expression of genes in these signatures between WT and *a2KO* embryos at E10.5 using the bRNAseq dataset. As expected, because *Hoxa2* is not expressed in PA1, we observe that the genes of the PA1 signature are highly expressed in both WT PA1 and *a2KO* PA1 (Fig. 5A). Interestingly, we do not see a noticeable increase in their expression in *a2KO* PA2 over WT PA2, which would be expected in a reversion of the mutant tissue to a PA1-like ground state. Equally strikingly, we see that many of the genes of the PA2 signature are likewise still strongly expressed in *a2KO* PA2 (Fig. 5B). The persistence of components of the PA2 signature and lack of induction of the PA1 signature further support the idea that there has not been a conversion of the transcriptional state of PA2 in *a2KO* embryos to a PA1-like state at this stage in association with the phenotypic transformation.

The expression of these transcriptional states is consistent between the bRNAseq and scRNAseq datasets. Aggregate gene expression of the PA1 signature is strongest in distal NCCs (cluster 3). We again observe strong expression of these genes in both WT and *a2KO* PA1, with limited expression in WT and *a2KO* PA2 (Fig. 5C). Aggregate gene expression of the PA2 signature is strongest in proximal NCCs (cluster 4). This expression is maintained in PA2 even in the absence of HOXA2 (Fig. 5D). Taken together, these data show that the *Hoxa2-/-* phenotypic reversion is not matched by a corresponding molecular transformation of PA2 to a PA1-like transcriptional ground state.

### Integration of transcriptomic datasets to identify putative HOXA2 targets in NCCs

To search for *Hoxa2*-dependent genes associated with axial specification, we identified genes that are differentially expressed between PA1 and PA2 in WT embryos, as well as those whose expression in PA2 changed in *a2KO* mutant embryos. For candidate genes specifying PA1 fate, we searched for genes that are expressed at higher levels in PA1 than PA2 of WT embryos, and also for those expressed at higher levels in PA2 of *a2KO* mutant embryos (potentially repressed by HOXA2) compared to WT. Conversely, for genes specifying PA2 fate, we identified genes that are expressed at higher levels in PA2 than PA1 of WT embryos, and those expressed at higher levels in PA2 of WT (potentially activated by HOXA2) compared to *a2KO* embryos. This two-step selection strategy identifies candidate genes that are downstream of *Hoxa2* and play potential roles in regulating the axial identity of NCCs.

To further exploit our transcriptomic datasets, we applied these criteria to both the bRNAseq dataset and the NCC and NCC-derived clusters (1-4) of the scRNAseq dataset, identifying 90 PA1 specifiers and 233 PA2 specifiers (Supplementary Tables S13 & S14). Among PA1 specifiers identified across datasets, we observe a very strong enrichment of ribosomal components and functions, including Gene Ontology (GO) terms for translation, peptide biosynthetic process, and RNA processing (Supplementary Table S15). The increase in expression of these components of protein synthesis in *a2KO* samples suggests they may be negatively regulated by *Hoxa2*. In contrast, PA2 specifiers are strongly enriched for transcription factors and transcriptional regulation, featuring GO terms such as protein binding, *cis*-regulatory region binding, and anatomical structure development (Supplementary Table S16). The decrease in expression levels of these genes in *a2KO* embryos indicates they may be positively regulated by HOXA2.

From candidate *Hoxa2*-responsive specifiers, some of the individual genes revealed by this comparative analysis offer insight into potential downstream pathways involved in the process (Table 2 and Fig. 6A-B). Among PA1 specifiers, we see a number of ribosomal subunits and rRNA processing genes, including *Rpl35, Rpl27, mt-Rnr2*, and *Npm1*. We also identified genes previously shown to be negatively regulated by *Hoxa2*, including *Lhx6, Alx4, Rspo2, Barx1*, and *Pitx1* (Bobola et al. 2003; Santagati et al. 2005; Kirilenko et al. 2011; Donaldson et al. 2012), validating the approach. Additionally, this analysis uncovered genes of interest that have not been previously characterized for their response to *Hoxa2* or their role in the specification of NCC axial identity, including *Tle4, Rnd3, Ift57*, and *Peg10*. Many of the candidate PA2 specifiers we identified are transcription factors, including well-characterized *Hoxa2* targets such as *Meis1, Meis2*, and *Meox1* (Kirilenko et al. 2011; Donaldson et al. 2012; Amin et al. 2015). In addition, some of the genes we identified as being dependent upon *Hoxa2*, are known to be functionally linked with specification of NCC, including *Zfp503, Zfp703*, and *Fzd4* (Donaldson et al. 2012). Furthermore, a number of novel targets emerged from this analysis, including *Ptn*, *Cntfr, Enho, Errfi1, 3ll0099E03Rik, Pou3f4, Hmx1*, and *Nr2f1*. Our selection strategy, based on analyses of bulk and scRNAseq datasets, has identified a list of candidate *Hoxa2*-responsive specifiers of NCC axial identity that include many novel and previously described genes involved in NCC development.

**Figure 6.**
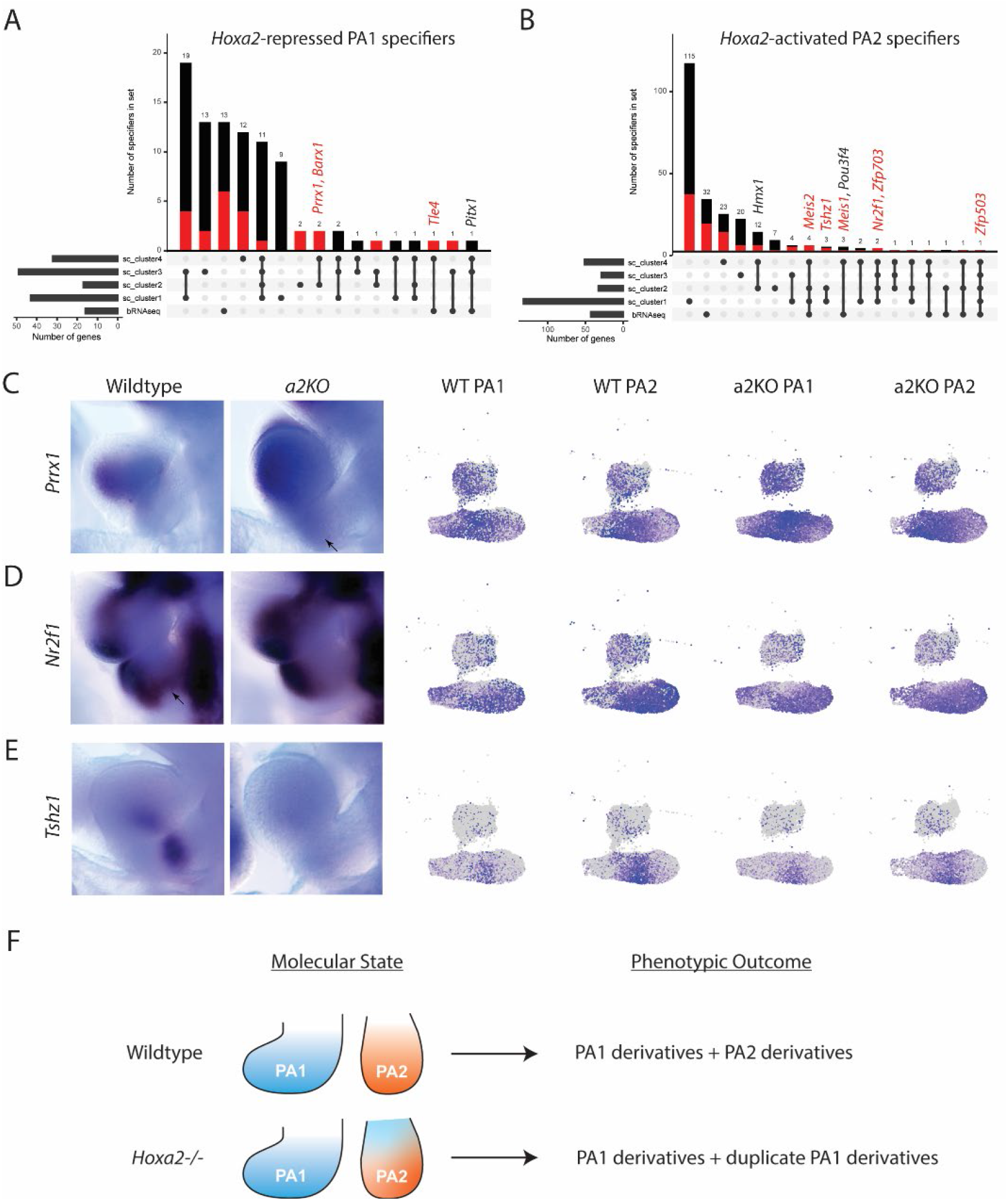
Putative axial specifiers show heterogeneous responses to Hoxa2 activity. A & B) UpSet plots depicting putative axial specifiers from bRNAseq and scRNAseq clusters 1-4. Red bars and text refer to putative direct targets of HOXA2 - those with nearby HOXA2 binding (Donaldson et al. 2012). A) *Hoxa2*-repressed PA1 specifiers. B) *Hoxa2*-activated PA2 specifiers. C-E) *In situ* hybridization and FeaturePlots showing expression in scRNAseq data for each gene in WT and *a2KO* embryos at E10.5, focusing on PA1 and PA2. C) PA1 specifier *Prrx1* shows expression in PA2 of *a2KO* embryos. PA2 specifiers *Nr2f1* (D) and *Tshz1* (E) show reduced expression in PA2 of *a2KO* embryos. F) Summary illustrating that the molecular state of PA2 does not correlate with the phenotypic outcome of PA2 in *a2KO* mutants.

**Table 2.** Putative *Hoxa2*-responsive axial specifiers identified through differential gene expression analysis from bRNAseq and scRNAseq NCC clusters.

| Hoxa2-repressed PA1 specifiers |  |  | Hoxa2-activated PA2 specifiers |  |  |
| --- | --- | --- | --- | --- | --- |
| *Ribosomal* | *Hoxa2-repressed* | *Novel* | *Hoxa2-activated* | *Novel* |  |
| Rpl35 | Lhx6 | Prrx1 | Meox1 | Pax9 | Enho |
| Rpl27 | Alx4 | Ift57 | Meis1 | Tbx15 | Errfi1 |
| mt-Rnr1 | Rspo2 | Rnd3 | Meis2 | Cntfr | Nr2f1 |
| mt-Rnr2 | Barx1 | Peg10 | Fzd4 | Pou3f4 | Hmx1 |
| Npm1 | Pitx1 | Tle4 | Zfp703 | Ptn | Tshz1 |

### Identification of putative direct targets of HOXA2 in axial specification of NCCs

To explore whether any of the candidate axial specifiers of PA1 and PA2 identified from our analyses may be directly regulated by *Hoxa2*, we leveraged published genome-wide binding data of endogenous HOXA2 protein in mouse PA2 at E11.5 (Donaldson et al. 2012). Despite the difference in developmental stages, this dataset presents an opportunity for comparison of our data at E10.5 with HOXA2 binding in differentiating NCCs at E11.5. In total, we find that 24/90 PA1 specifiers (26.7%) and 91/233 PA2 specifiers (39.1%) have at least one nearby HOXA2 binding peak, suggesting possible direct regulation by HOXA2 (bold in Table 2 and Fig. 6A-B). In this comparison we observe that a number of genes that have been previously identified as direct targets of HOXA2 show nearby binding, including putative PA1 specifiers *Barx1* and *Rspo2*, as well as putative PA2 specifiers *Fzd4, Meis1*, and *Meis2*. However, other published targets of *Hoxa2* may be indirect, as evidenced by a lack of nearby HOXA2 binding. These include putative PA1 specifiers *Pitx1* and *Alx4*, as well as putative PA2 specifier *Meox1*.

In conclusion, this integration of DNA binding data with our analysis has allowed us to identify candidate axial specifiers of NCC identity which are directly and indirectly downstream of *Hoxa2* in patterning the PAs. Together, these data provide insight into the molecular mechanisms and cell populations through which HOXA2 establishes the axial identity of PA2.

### Putative axial specifiers show diverse responses to HOXA2 activity

To deepen our understanding of the mechanisms by which putative axial specifiers uncovered through our analyses of bRNAseq and scRNAseq data impart axial identity, we sought to validate our findings and explore how these genes behave across PA1 and PA2 in both WT and *a2KO* embryos. We observe expression of these genes throughout the NCCs and a diverse set of responses to the absence of HOXA2. *Prrx1*, a putative PA1 specifier, has been shown to be essential for skeletogenesis, including correct specification of the middle ear (Martin et al. 1995) but had not previously been associated with *Hox* gene expression. We observe expression of *Prrx1* in PA2 of *a2KO* embryos, but not their WT littermates (Fig. 6C). Putative PA2 specifiers *Nr2f1* and *Tshz1* show the converse trend - reduced expression in *a2KO* PA2 when compared to WT littermates (Fig. 6D-E). *Nr2f1* has been implicated in positive regulation of NCC enhancers (Rada-Iglesias et al. 2012) and the *Nr2f* family of nuclear receptors has been shown to play a role in the specification of bone and cartilage in the zebrafish jaw (Barske et al. 2018). *Tshz1* has been shown to play a role in middle ear development (Coré et al. 2007). However, neither gene has previously been shown to interact with *Hox* genes.

The genes identified by the comparative analyses above show unique spatial expression patterns and responses to HOXA2 activity, suggesting distinct roles for these genes in *Hoxa2-* mediated axial specification. The *in vivo* observations also help to validate the differential association of these putative targets with specific subsets of NCCs in clusters 1-4 uncovered by our datasets (Fig. 6A-B and Supplementary Tables S13 and S14). We believe that many other genes identified in our comparative analysis share similar properties and with thorough functional validation will enhance our understanding of *Hox*-mediated NCC axial specification. In contrast to what might be expected based on the observed phenotypic transformation in *Hoxa2* mutants, we conclude that there is not a singular molecular ground state in NCCs that is dramatically switched by HOXA2 to impart a PA2 fate. Rather, we see different transcriptional profiles for PA1 and PA2 with subsets of NCCs expressing distinct sets of genes in each PA. Moreover, we see that many of the putative axial specifiers identified here do not depend upon HOXA2 in all NCCs but rather show marked differences in subsets of cells. This suggests that subsets of cells are likely to be responsible for imparting specific cell fates and morphogenic features in the PAs (Fig. 6F). This context dependent role for HOXA2 uncovers complexity in our understanding of NCC axial specification and highlights an unexplored heterogeneity of the system.

## Discussion

*Hox* genes play a key role in craniofacial development by specifying the axial identity of neural crest cells migrating into the pharyngeal arches of the developing head. In this study, we have used genomic approaches to investigate the transcriptional programs that underlie the role of *Hoxa2* in specifying the fate of PA2 NCCs. *Hoxa2* is believed to be a selector gene that serves as a regulatory switch to convert a PA1 ground state into a unique PA2 identity. Comparing the bulk and single cell transcriptomes of PA1 and PA2 in wildtype and *Hoxa2* mutant embryos during NCC migration and differentiation, we find that the phenotypic transformations observed in *Hoxa2* mutants is not matched by a corresponding molecular transformation of PA2 to a PA1-like transcriptional ground state. This separation of phenotypic and molecular states has significant implications for our understanding of NCC biology and craniofacial development. The scRNAseq analyses also reveal heterogenous expression patterns in NCC populations within the PAs and the changes observed upon loss of HOXA2 suggest that different subsets of NCCs may respond to HOXA2 activity in distinct manners related to their ultimate fate. Our findings raise a number of interesting issues important for understanding axial specification of NCCs and the role of HOXA2 in this process.

### Transcriptional analysis suggests that there is not a molecular NCC ground state

Previous work has demonstrated the existence of a phenotypic ‘ground state’ among NCCs colonizing the PAs in the developing mouse embryo, which is modified by *Hox* gene expression to produce axial-specific derivatives (Minoux et al. 2009). In this study, we set out to understand the molecular underpinnings of this phenotypic ground state and identify the mechanisms by which *Hox* genes impart axial identity. To address this question, we began by comparing WT and *a2KO* PA1 and PA2 transcriptional profiles throughout NCC migration and differentiation (E9.0-E10.5). In WT embryos, we observe large differences in gene expression profiles between PA1 and PA2 from the earliest point in our timecourse - E9.0, when NCCs are migrating into the PAs (Fig. 2A). Hence, NCCs migrating into different PAs do not appear to emerge from the neural tube with a common or shared transcriptional signature that is progressively modified to generate unique axial identities. This contrasts with what might be expected from populations of cells known to maintain plasticity and be highly responsive to environmental signals as they migrate at E9.0 (Trainor and Krumlauf 2000a; Trainor and Krumlauf 2000b; Schilling et al. 2001). Because the cells entering the PAs have distinct gene expression patterns based on their axial level of origin prior to delamination (Hindbrain *Hox* code in Fig. 1B), and have been previously shown to be primed to respond to different environmental cues in the PAs (Trainor and Krumlauf 2001; Trainor et al. 2002), it is not unreasonable to expect their gene expression profiles will differ during early phases of migration.

In comparing the transcriptomes of PA1 and PA2 in both WT and *a2KO* embryos, the existence of a molecular ground state underlying the previously described phenotypic transformation of PA2 to a PA1 identity would suggest that the transcriptome of PA2 reverts to a PA1-like transcriptome in *a2KO* embryos, mirroring the phenotypic reversion. We did not observe such a transformation or switch in the molecular identity of NCCs in PA2 over the course of their migration and differentiation (Fig. 2C). This argues against the existence of a common PA1-like molecular ground state in NCC development that is modified by *Hox* expression. The comparison of WT PA1 and *a2KO* PA1 also revealed a surprising number of differentially expressed genes (Supplementary Fig. S5 and Supplementary Tables S17-21). This is unexpected, as *Hoxa2* is not thought to be expressed within the NCCs giving rise to PA1 derivates (Couly et al. 1998) and no phenotypes in PA1 structures have been observed in *Hoxa2* mutant embryos (Gendron-Maguire et al. 1993; Rijli et al. 1993). These differences in the molecular character of PA1 generated by the loss of *Hoxa2* may reflect an unexpected early role for the gene in modulating properties of NCCs in the hindbrain before they emigrate to colonize PA1. Alternatively, the differences could be a consequence of altered signaling between PAs in mutant embryos or reflect previously undetected expression of *Hoxa2* in PA1 NCCs. In this regard, it is worth noting that our scRNAseq data suggest the presence of a small number of *Hoxa2* expressing cells in PA1 (Supplementary Fig. S4). Regardless of the underlying mechanism, this unexpected effect on gene expression in PA1 upon loss of HOXA2, reveals complexities in the establishment of the PA1 transcriptional signature that are not consistent with a molecular ground state model.

In considering alternatives to a ground state model, one concern in studying a heterogeneous sample, such as the PA, using bRNAseq is the noise inherent in the system, with gene expression changes potentially amplified or lost for our tissue of interest (NCCs) because of reads from another tissue. It is possible that a small set of cells, or a small set of genes are responsible for the phenotypic transformation observed in *a2KO* PA2, which would not shift the general transcriptome of the entire PA. Consistent with this idea, previous work indicates that different cells may respond to *Hoxa2* expression in different ways - with distinct requirements for levels and timing of *Hoxa2* expression within NCCs (Ohnemus et al. 2001; Santagati et al. 2005).

### scRNAseq reveals heterogeneity of differentiating NCC populations in the mouse embryo

In recent years, scRNAseq experiments have been used to shed light on the dynamics of embryonic development, from whole-embryo studies (Cao et al. 2019; Soldatov et al. 2019) to in-depth characterization of particular developmental processes (Xu et al. 2019; Tambalo et al. 2020). Here, we took advantage of this approach to explore the heterogeneity of the PAs and characterize the role of HOXA2 in imparting PA2 NCC identity at single-cell resolution. We focused our analysis on E10.5, the start of NCC differentiation, due to the strong effects of HOXA2 observed in our bRNAseq analysis and previous work characterizing HOXA2 activity at this timepoint (Santagati et al. 2005; Donaldson et al. 2012; Amin et al. 2015). We generated transcriptional profiles for 67,674 cells across our four samples (PA1 and PA2 from WT and *a2KO* embryos), which aligned into 11 clusters, four of which correspond to NCCs. We focused primarily on uncommitted NCCs (cluster 1) and NCC-derived bone and cartilage (clusters 2-4), which together comprise 80.2% of the total cells isolated.

Our data show strong patterns in the expression of both proximal-distal and rostral-caudal markers in clusters 2-4 consistent with previously published observations (Xu et al. 2019). This suggests that the clustering, a reflection of transcriptional similarity between cells, is informed by the location of NCCs within the PA (Fig. 4A-B). Moreover, the expression of chondrogenic markers *Sox9* and *Col2a1* is similar between WT PA1 and PA2, while osteogenic markers *Msx1* and *Msx2* are enriched in PA1 (Fig. 4C), consistent with the formation of cartilage in both tissues but bone primarily in PA1 (Dash and Trainor 2020). This trend is consistent in *a2KO* embryos (Supplementary Fig. S3), implying that intramembranous ossification is not increased in *a2KO* PA2 at this point in development, although it likely is at a later point as duplicate PA1 bone structures are being formed in the arch.

Consistent with previous observations, we see *Hoxa2* expression throughout PA2, expressed primarily in NCCs and NCC-derived clusters (Fig. 4D-F). It is important to note that because of the structure of the *a2KO* mutant allele, *Hoxa2* transcript is produced in the homozygous null embryos, but not translated. This serves as a lineage tracer for *Hoxa2-* expressing cells and shows they are not lost in *a2KO* embryos, demonstrating that HOXA2 is not required for their survival. Interestingly, we do not observe strict segregation of *Hoxa2*-expressing cells and *Sox9*-expressing cells (Kanzler et al. 1998). This is potentially due to the collection of these cells early in the differentiation process, a pattern that is refined over the course of developmental time. The higher levels of *Hoxa2* expression at the proximal end of the NCC-derived bone/cartilage clusters are also consistent with previous observations that the proximo-caudal portions of PA2 require higher levels of HOXA2 for proper skeletal development (Ohnemus et al. 2001).

### Behavior of PA-specific transcriptional signatures in WT and *a2KO* embryos indicates lack of a molecular NCC ground state

Based on bRNAseq transcriptomes of PA1 and PA2 in WT embryos at E10.5, we utilized differential expression analysis to identify a set of genes enriched in PA1, referred to as the PA1 transcriptional signature, and a set of genes enriched in PA2, referred to as the PA2 transcriptional signature. By establishing a reference set of genes characteristic of each PA in WT embryos, we provided a framework for comparison to determine how tissue identity changes in the absence of HOXA2. We also used this as a reference to shed light on the molecular mechanisms of HOXA2 activity and characterize the molecular underpinnings of the *Hox*-free PA1 phenotypic ground state and how it is altered in *Hoxa2^-/-^* embryos.

Based on the phenotypic transformation of PA2 NCC derivatives to duplicate PA1 structures in the absence of HOXA2 (Minoux et al. 2009), we expect an accompanying transformation of the *a2KO* PA2 transcriptome to a PA1-like molecular ground state. However, the PA1 and PA2 signatures in both our bRNAseq and scRNAseq datasets revealed a retention of the general PA2 signature and a lack of upregulation of PA1 signature genes in PA2 of *a2KO*, embryos (Fig. 5). These data strongly suggest the absence of a true molecular NCC ground state that is globally altered by *Hoxa2* expression in PA2 NCCs. The resolution afforded by scRNAseq in this context enables us to identify transcriptional changes regardless of potential tissue heterogeneity and noise and illustrates its value in evaluating models surrounding the molecular basis of developmental and evolutionary phenotypes.

Our observations are consistent with a published comparison of mouse NCCs at E10.5 from WT PA1, WT PA2, and PA1 with ectopic *Hoxa2* expression (Minoux et al. 2017). In that study, *Hoxa2* expression alters the PA1 transcriptional profile, but *Hoxa2*-expressing PA1 NCCs do not cluster with WT PA2 cells, as would be expected if HOXA2 was sufficient to drive a molecular transformation corresponding to the phenotypic one (Kitazawa et al. 2015). The effects of HOXA2 in PA2 specification appear to be more subtle than expected, giving a glimpse into what appears to be a complex regulatory picture consisting of cross-regulation with other *Hox* genes (Tümpel et al. 2007), cofactors such as *Pbx* and *Meis* (Amin et al. 2015; Parker et al. 2019), axial signaling programs such as FGF (Trainor et al. 2002), and epigenetic regulation (Minoux et al. 2017). Although the effect of *Hoxa2* knockout (Fig. 2C & Fig. 5) or overexpression (Minoux et al. 2017) on the average transcriptome of NCCs is limited in scope, it nevertheless results in a dramatic phenotypic transformation. Our scRNAseq dataset reveals that subsets of *a2KO* PA2 NCCs do show reduction in expression of PA2 signature genes and upregulation of PA1 signature genes, suggesting that a molecular transformation may be occurring on a limited scale. This prompted us to further investigate the heterogeneity of NCCs and their response to HOXA2.

### HOXA2 imparts PA2 identity in subsets of NCCs

Several studies have now looked at transcriptional differences between PA1 and PA2 in mouse embryos at various stages, seeking to identify key characteristics and components in establishment of axial identity (Brunskill et al. 2014; Lumb et al. 2017; Minoux et al. 2017). Other studies have identified a small set of genes regulated by *Hoxa2* within NCCs (Kanzler et al. 1998; Santagati et al. 2005; Kirilenko et al. 2011; Donaldson et al. 2012; Minoux et al. 2013). Despite these extensive efforts and the known role of HOXA2 in specifying NCC axial identity, the link between HOXA2 activity and PA2 fate has not been well characterized at this point. Here, we identified *Hoxa2*-responsive axial specifiers for both PA1 and PA2. In addition to finding a number of novel candidates, several of the genes we identified have been previously shown to have an axial-specific bias and others to respond to HOXA2 activity. Our analyses describing putative *Hoxa2*-responsive axial specifiers establish new regulatory links between these genes and the regulatory network governing the specification of NCC identity. This provides functional and mechanistic insights into the role of HOXA2 in the establishment of NCC axial identity.

PA1 specifiers are strongly enriched for translation and related processes (Supplementary Table S15). There is increasing evidence from studies on craniofacial abnormalities in humans and vertebrate model systems that it is important to maintain the proper balance of ribosome biogenesis in cranial NCCs (Dixon et al. 2006; Jones et al. 2008; Weaver et al. 2015; Terrazas et al. 2017; Watt et al. 2018). Moreover, individual genes such as *Npm1* that emerge as putative HOXA2-repressed PA1 specifiers have been implicated in processes including epithelial-to-mesenchymal transition, such as that observed when NCCs delaminate from the neural tube to begin migration (Prakash et al. 2019). In contrast, PA2 specifiers are largely enriched for developmental processes and morphogenesis, with many of the identified genes acting as transcription factors (Supplementary Table S16). It is encouraging that we identify genes previously implicated in NCC-related processes, such as *Nr2f1* (Fig. 6D), as well as genes shown to play a role in middle ear development, such as *Tshz1* (Fig. 6E), which have not previously been connected to *Hox* gene activity. These connections enable us to identify specific developmental processes through which *Hoxa2* likely acts to impart PA2 identity.

The differences observed between specifiers of PA1 and PA2 likely speak to the processes occurring at different axial levels at this time. The enrichment for ribosomal components in PA1 may suggest higher levels of proliferation in this tissue, a hypothesis supported by the fact that the skeletal structures derived from PA1 NCCs are significantly larger than those in PA2, thus requiring a greater expansion of the NCC population. Moreover, there may be differences in developmental timing between PA1 and PA2, with PA1 forming earlier than PA2, resulting in the gene expression differences observed in our data.

The advent of genomic tools to address questions in developmental biology marks a shift in our approach to characterizing phenotypes and the activity of individual genes. These data provide opportunities to explore the correlation, or lack thereof, between phenotypic effects and the underlying transcriptional differences that lead to them. Studies are beginning to emerge that draw a distinction between morphological and molecular phenotypes (Dooley et al. 2019). Likewise, here we show that the striking phenotypic ground state of mouse NCCs is nonetheless not accompanied by a corresponding molecular ground state (Fig. 6F). In light of the differences we observe in transcriptional regulation between subsets of NCCs, it would be invaluable to determine whether there are corresponding changes in HOXA2 occupancy within the genome that correlate with them. Though we integrated a previously published dataset of HOXA2 binding in PA2 (Donaldson et al. 2012), the rapid development of low-input, high-resolution methods to assay transcription factor binding such as Cut & Run (Hainer and Fazzio 2019) and ChIPmentation (Schmidl et al. 2015) will likely prove a meaningful step toward identifying such correlations at higher resolution and with more confidence. Furthermore, recent data show strong epigenetic differences between craniofacial tissues across axial levels (Minoux et al. 2017). The relationships between these *cis*-regulatory features, HOXA2 binding, and the transcriptional status of subsets of NCCs will provide a wealth of data toward understanding axial-specific processes.

## Materials and Methods

### Mouse husbandry and embryo collection

All mouse work was performed in the Laboratory Animal Services Facility at the Stowers Institute for Medical Research under IACUC approved protocols RK-2016-0164 and RK-2019-094. Euthanasia procedures were performed in accordance with recommendations by the American Veterinary Medical Association. All mice used in this study were maintained on a CBA/Ca/J x C57BL/10 background.

### CRISPR-Cas9 mutation to create Hoxa2-ATG-XhoI mouse line

A CRISPR-Cas9 gene editing approach was developed to mutate a 7 bp region including part of the start codon of HOXA2 (5’-AGGCC**ATG-3’**), by deleting 2 bp and converting it into an XhoI site (5’-CTCGA**G-3’**) - hereafter referred to as the *a2KO* allele. A chimeric guide RNA was designed to target the start codon of HOXA2, consisting of the oligos 5’-CACCGCCGAGGGGGCTCCAAGGAGA-3’ and 5’-AAACTCTCCTTGGAGCCCCCTCGGC-3’, which were annealed and ligated into the pX330 plasmid (Cong et al. 2013). Together with a homology oligo (5’-CTTGCCCCCCCAAAGCCCCTCCAAAAGAGGGAACTTTTCCTCCGAGGGGGCTCCAAGGAGA**CTCGAG**AA TTACGAATTTGAGCGAGAGATTGGTTTTATCAATAGCCAGCCGTCGCTCGCTGAGTGC-3’), the pX330 plasmid was microinjected into one-cell CBA/Ca/J x C57BL/10 embryos collected from superovulated donor mice. The next day, surviving two-cell embryos were transferred into pseudopregnant CBA/Ca/J x C57BL/10 females and genotyped upon weaning. Founders were maintained on this background.

### Bulk RNAseq preparation and sequencing

Embryos were collected from heterozygous *a2KO* matings at approximately E9.0, E9.5, E10.0, and E10.5, with the day of identification of a vaginal plug defined as E0.5. The number of somite pairs were counted to more accurately stage each embryo and their yolk sacs used to determine genotype. To minimize variation in staging between individual embryos, we narrowly defined the number of somites appropriate for each stage (E9.0: 16-17 somites, E9.5: 22-24 somites, E10.0: 28-29 somites, E10.5: 35-36 somites) and only used wildtype (WT) or homozygous *a2KO* mutant embryos falling within the respective ranges. PA1 and PA2 were individually isolated from each embryo by manual dissection in ice-cold phosphate-buffered saline (PBS), flash frozen in liquid nitrogen, and then stored at −80 °C. Following genotyping, Ambion TRIzol (catalog number 15596026) was added to each frozen sample, vortexed to homogenize, and the Zymo Research Direct-zol mini-prep kit (catalog number R2052) with on-column DNase treatment was used to extract RNA. This was performed for PA1 and PA2 of individual embryos, resulting in 3-5 WT and *a2KO* biological replicates at each developmental timepoint.

RNA quantification and quality control were performed using an Agilent 2100 Bioanalyzer. All samples had RIN scores >9.0. The Takara Clontech SMART-seq v4 ultra low input RNA kit (catalog number 634891), Illumina Nextera XT Library prep kit (catalog number FC-131-1096), and Illumina Nextera XT Index kit (catalog number FC-131-2001) were used for polyA-selected cDNA preparation and library construction. Library quality was checked using an Agilent 2100 Bioanalyzer, then pooled and loaded onto five lanes of an Illumina HiSeq flow cell to sequence 50 bp single reads for a total of 20-30X genomic coverage.

### Bulk RNAseq data analysis

Reads were aligned to mm10 (Ensembl 91) using Tophat 2.1.1 (Kim et al. 2013). Downstream analysis was performed in R 3.3.2 using EdgeR quasi-likelihood pipeline 1.4.1 (Chen et al. 2016) for differential expression analysis with default settings. Differentially expressed genes between samples were called with adjusted p < 0.05.

### Single-cell RNAseq preparation and sequencing

Homozygous *a2KO* mutant embryos do not display obvious morphological phenotypes at E10.5 that would enable them to be distinguishable from wildtype or heterozygous littermates. Hence, it was necessary to develop a rapid genotyping method to identify wildtype and homozygous *a2KO* embryos which is also compatible with subsequent processing steps for scRNAseq experiments. Mouse embryos were collected with their extra-embryonic membranes intact at E10.5 from *a2KO* heterozygous matings and kept in Tyrode’s Solution (8.0g NaCl, 0.2g KCl, 0.2g CaCl2, 0.21g MgCl_2_*6H2O, 0.57g NaH2PO4*H2O, 1.0g glucose, 1.0g NaHCO3 per 1L DEPC treated H2O, pH 7.4) at room temperature until ready to proceed with dissections of the PAs. Approximately 1 mm^2^ piece of the yolk sac was collected for each embryo and genotyped using ThermoFisher Phire Tissue Direct PCR Master Mix Kit (catalog number F170S) to rapidly identify homozygous *a2KO* embryos. WT embryos were collected at E10.5 from CBA/Ca/J x C57BL/10 matings.

PAs from homozygous *a2KO* (n = 5) and WT (n = 6) embryos were manually dissected in ice-cold DEPC-treated phosphate-buffered saline (DPBS, Sigma catalog number D8537). Individual PAs were pooled into four samples (WT PA1, WT PA2, a2KO PA1, a2KO PA2) and pooled samples dissociated by incubation in 0.25% Trypsin + EDTA (Gibco catalog number 25200-056) for 3 minutes with two rounds of manual disruption by pipetting. The reaction was stopped with fetal bovine serum and samples were washed two times in DPBS. Single-cell suspensions were loaded on a Nexelome Cellometer Auto T4 to assess cell count and viability. All samples had >90% viability and were loaded on a 10X Chromium Single Cell Controller. Libraries were prepared using the Chromium Single Cell 3’ Reagents Kits v3 (CG000183 Rev A). Library quality was checked on an Agilent 2100 Bioanalyzer, then pooled and loaded onto two Illumina NovaSeq S1 flow cells to sequence paired reads consisting of: 28 bp cell barcode & UMI, 8 bp i7 index, and 91 bp read. All samples were sequenced to a read depth >60,000 reads/cell.

### Single-cell RNAseq data analysis

Reads were aligned to mm10 (Ensembl 98) using CellRanger 3.0.0. Downstream analysis was performed in R 3.5.2 using Seurat 3.1.0 (Butler et al. 2018; Stuart et al. 2019). Cells were removed from analysis if they had mitochondrial percentage greater than 5% or fewer than 200 genes expressed. DoubletFinder 2.0 (McGinnis et al. 2019) was used to remove ~20% of cells identified as doublets (based on extrapolation of 10X Chromium reported multiplets for a given number of recovered cells). After this final filtering step, a total of 67,674 cells remained for downstream analysis. Further analysis was performed on these cells using the standard Seurat Integration pipeline, with 2000 variable features and 40 principal components used for uniform manifold approximation and projection (UMAP) and clustering (at resolution = 0.5). Cluster identity was assigned based on marker genes identified using FindAllMarkers() function with a minimum of 10% of cells expressing a gene and log fold change greater than 0.25, using marker genes annotated in (Cao et al. 2019). Exploration of cell spatial distribution was performed via analysis of genes described in (Xu et al. 2019). To identify genes differentially expressed between samples, the FindMarkers() function was used on a cluster-by-cluster basis to make pairwise comparisons between samples, with a minimum of 10% of cells expressing a gene and log fold change greater than 0.10.

### Gene ontology and pathway analysis

For lists of genes of interest, gProfiler g:GOSt functional enrichment analysis was used to obtain GO, KEGG, and other functional characterization in R3.6.1 using gprofiler2 version 0.1.9 with a statistical significance threshold of p < 0.05. Further information on gProfiler methodology is described in (Raudvere et al. 2019).

### In situ hybridization

Mouse embryos homozygous for the *a2KO* mutation and WT littermates were collected at E10.5 and fixed overnight in 4% paraformaldehyde in PBS. Probe generation and hybridization performed as previously described (Ariza-McNaughton and Krumlauf 2002). Primers used to generate probes for *Hoxa2* targets from cDNA as follows:

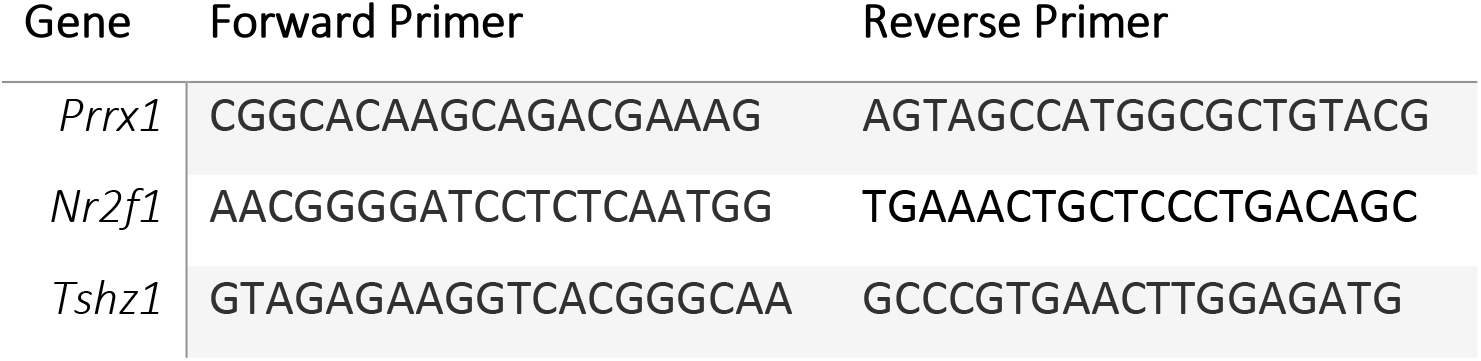

Each gene was assayed in 4-10 embryos of each genotype and only those with consistent patterns were considered significant. Z-stack images were acquired with a Leica MZ16 steromicroscope and subsequently processed with Helicon Focus 6.8.0 software (Helicon Soft Ltd.) to create a projection of all imagines into a single high-focus photo.

### Skeletal staining

Mouse embryos heterozygous or homozygous for the *a2KO* allele along with WT littermates were collected at E18.5, euthanized in PBS on ice for 1 hour, and then fixed and stored in ice cold 100% ethanol. Staining with Alcian Blue and Alizarin Red was performed as previously described (Rigueur and Lyons 2014).

## Data Availability

The bulk and single-cell RNAseq data generated as part of this manuscript have been deposited into the National Center for Biotechnology Information Gene Expression Omnibus (NCBI GEO) database - accession number GSE164111. Files generated during analysis have been included as Supplementary Tables where deemed appropriate. Original data underlying this manuscript can be accessed from the Stowers Original Data Repository at http://www.stowers.org/research/publications/libpb-1598.

## Supplementary Table Legends

<u>Table S1.</u> Summary table of genes differentially expressed in bRNAseq data between WT PA1 and WT PA2 across timepoints.

<u>Table S2.</u> Results of differential expression analysis from bRNAseq data between WT PA1 and WT PA2 at E9.0 including adjusted p-value and log(fold change).

<u>Table S3.</u> Results of differential expression analysis from bRNAseq data between WT PA1 and WT PA2 at E9.5 including adjusted p-value and log(fold change).

<u>Table S4.</u> Results of differential expression analysis from bRNAseq data between WT PA1 and WT PA2 at E10.0 including adjusted p-value and log(fold change).

<u>Table S5.</u> Results of differential expression analysis from bRNAseq data between WT PA1 and WT PA2 at E10.5 including adjusted p-value and log(fold change).

<u>Table S6.</u> Summary table of genes differentially expressed in bRNAseq data between WT PA2 and *a2KO* PA2 across timepoints.

<u>Table S7.</u> Results of differential expression analysis from bRNAseq data between WT PA2 and *a2KO* PA2 at E9.0 including adjusted p-value and log(fold change).

<u>Table S8.</u> Results of differential expression analysis from bRNAseq data between WT PA2 and *a2KO* PA2 at E9.5 including adjusted p-value and log(fold change).

<u>Table S9.</u> Results of differential expression analysis from bRNAseq data between WT PA2 and *a2KO* PA2 at E10.0 including adjusted p-value and log(fold change).

<u>Table S10.</u> Results of differential expression analysis from bRNAseq data between WT PA2 and *a2KO* PA2 at E10.5 including adjusted p-value and log(fold change).

<u>Table S11.</u> Marker genes for all clusters in scRNAseq dataset at E10.5.

<u>Table S12.</u> Genes corresponding to PA1 and PA2 transcriptional signatures based on WT PA1 vs. WT PA2 differential expression analysis.

<u>Table S13.</u> Putative *Hoxa2*-repressed PA1 specifiers. Genes identified from bRNAseq, as well as scRNAseq clusters 1-4. For each gene, 1 corresponds to being statistically significant in a given dataset, 0 indicates not significant. In the last column, B indicates it is the nearest gene to a HOXA2 binding site from (Donaldson et al. 2012), N indicates that it is not.

<u>Table S14.</u> Putative *Hoxa2*-activated PA2 specifiers. Genes identified from bRNAseq, as well as scRNAseq clusters 1-4. For each gene, 1 corresponds to being statistically significant in a given dataset, 0 indicates not significant. In the last column, B indicates it is the nearest gene to a HOXA2 binding site from (Donaldson et al. 2012), N indicates that it is not.

<u>Table S15.</u> GO terms for putative *Hoxa2*-repressed PA1 specifiers.

<u>Table S16.</u> GO terms for putative *Hoxa2*-activated PA2 specifiers.

<u>Table S17.</u> Summary table of genes differentially expressed in bRNAseq data between WT PA1 and *a2KO* PA1 across timepoints.

<u>Table S18.</u> Results of differential expression analysis from bRNAseq data between WT PA1 and *a2KO* PA1 at E9.0 including adjusted p-value and log(fold change).

<u>Table S19.</u> Results of differential expression analysis from bRNAseq data between WT PA1 and *a2KO* PA1 at E9.5 including adjusted p-value and log(fold change).

<u>Table S20.</u> Results of differential expression analysis from bRNAseq data between WT PA1 and *a2KO* PA1 at E10.0 including adjusted p-value and log(fold change).

<u>Table S21.</u> Results of differential expression analysis from bRNAseq data between WT PA1 and *a2KO* PA1 at E10.5 including adjusted p-value and log(fold change).

## Acknowledgements

We are grateful to members of the Krumlauf lab for extensive discussion of this work, especially Christof Nolte who assisted with generation of the *a2KO* mice. We are also grateful to William Munoz for assistance with imaging *in situ* hybridization in embryos. We appreciate the assistance of Jason Morrison and Paul Kulesa in getting single-cell RNAseq working in our system. We thank Heidi Monin, Kathleen Zapien, and other members of the Laboratory Animal Services staff at SIMR for maintaining and caring for the mice used in this study. We thank Allison Peak and other members of the Sequencing and Discovery Genomics facility for 10X Chromium setup and utilization, library preparation, and sequencing services. We thank Madelaine Gogol and Chris Seidel for support and assistance with data analysis as well as the Computational Genomics core for preliminary data preparation. We are grateful to Mark Miller for illustration and figure preparation. We would also like to acknowledge the University of Kansas Medical Center Genomics Core for sequencing support, as well as the *Kansas Intellectual and Developmental Disabilities Research Center (NIH U54 HD 090216), the Molecular Regulation of Cell Development and Differentiation – COBRE (P30 GM122731-03) and the NIH S10 High-End Instrumentation Grant (NIH S10OD021743)* at the University of Kansas Medical Center, Kansas City, KS 66160. Research reported in this publication was supported by the National Institute of Dental & Craniofacial Research of the National Institutes of Health under Award Number F31DE028469 to IP. The content is solely the responsibility of the authors and does not necessarily represent the official views of the National Institutes of Health. This research is also supported by grants from the Stowers Institute for Medical Research to RK (#1001) and PAT (#1008). This work was performed to fulfill, in part, requirements for IP’s thesis research in the Graduate School of the Stowers Institute for Medical Research.

---

**Supplementary Figure S1.**
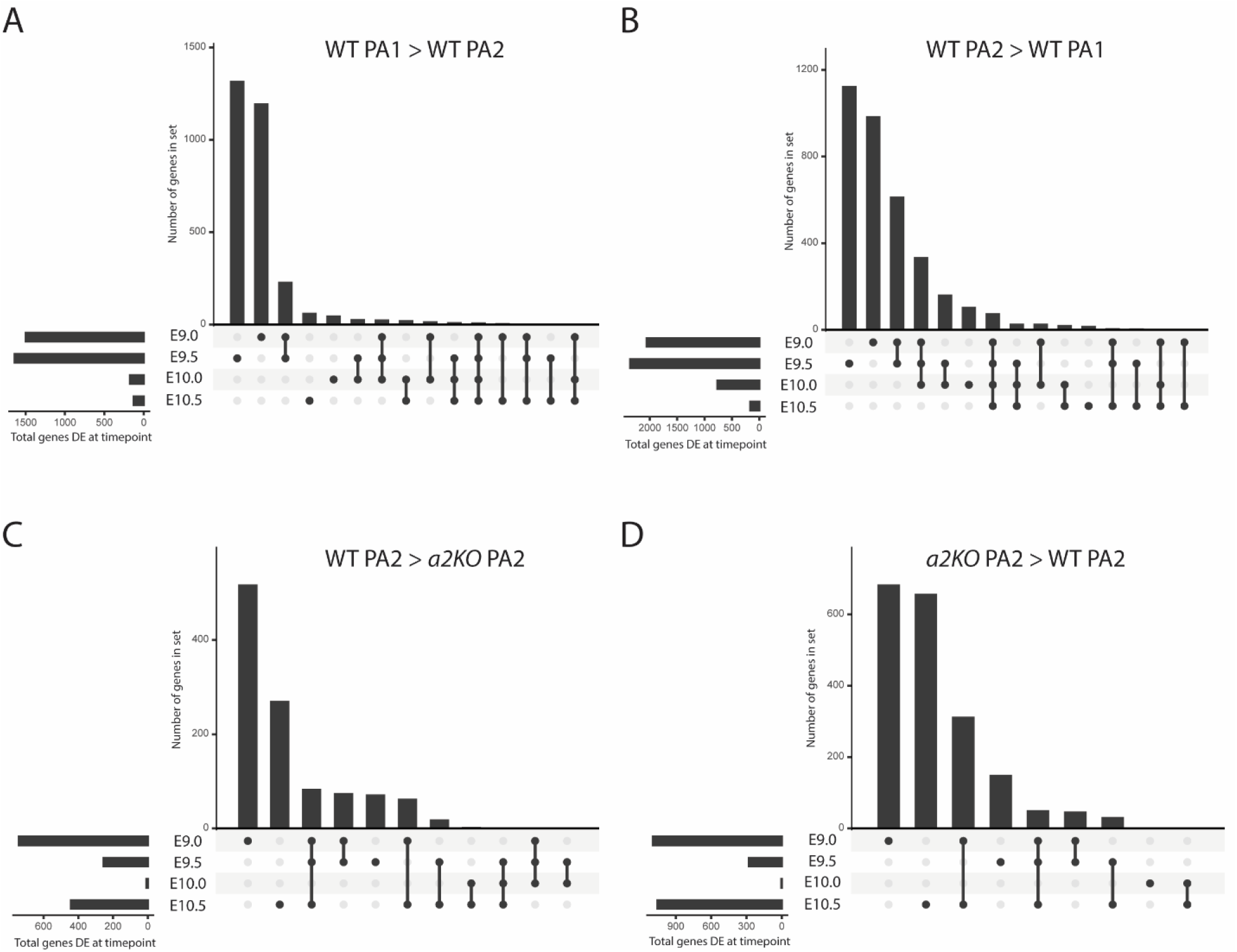
UpSet plots depicting differentially expressed genes across timepoints in bRNAseq dataset. UpSet plots depict overlapping data in a clearer format than Venn diagrams. Each timepoint is listed at the left, with dots (connected by lines) showing the datasets that a given group of genes (bar) belongs to. For further information on UpSet plots, refer to (Lex et al. 2014). A) Genes with higher expression in WT PA1 than WT PA2 at any timepoint. B) Genes with higher expression in WT PA2 than WT PA1 at any timepoint. C) Genes downregulated in PA2 in the absence of HOXA2. D) Genes upregulated in PA2 in the absence of HOXA2.

**Supplementary Figure S2.**
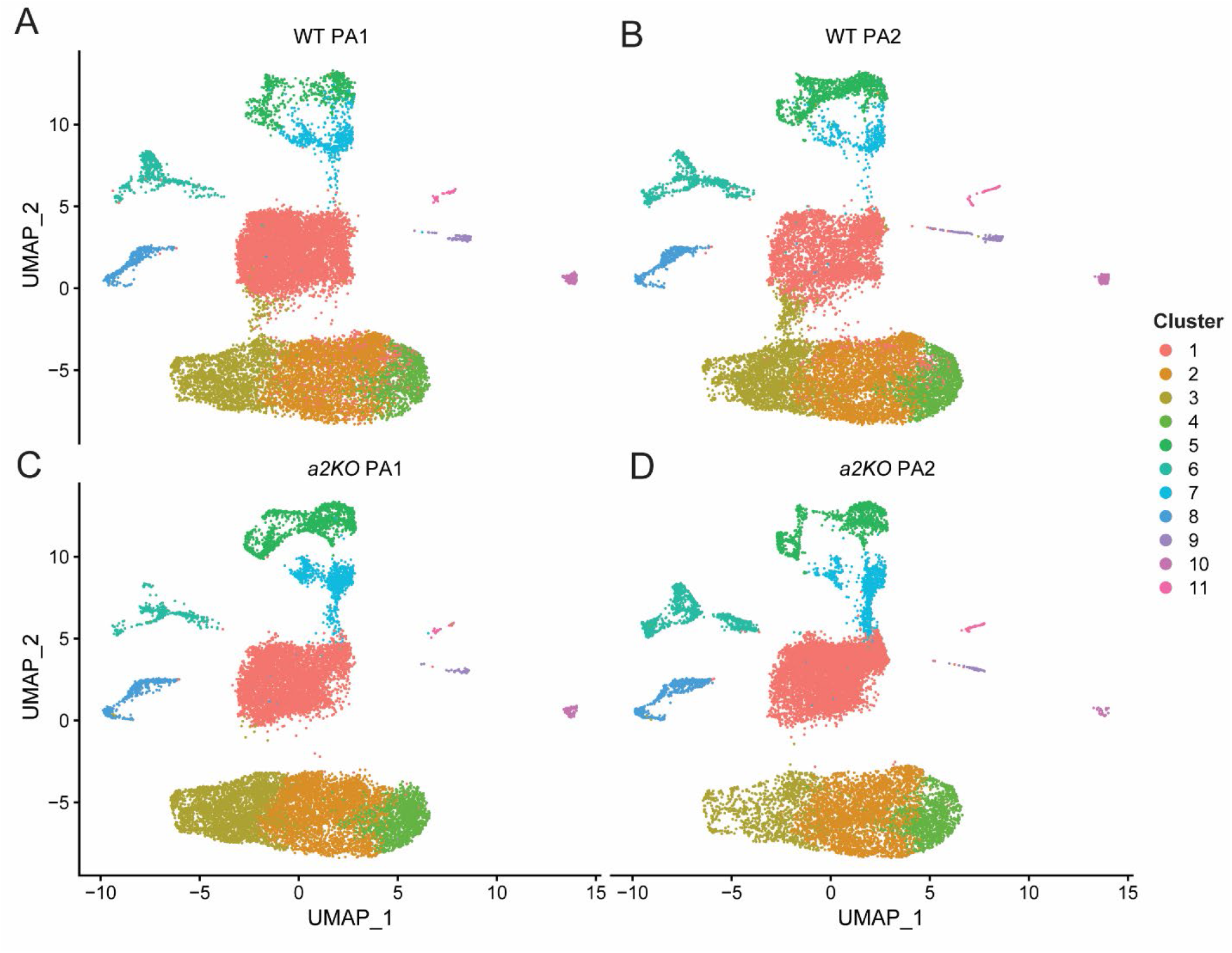
UMAP visualization of cells from each of the four samples. Cells colored by cluster as defined in Figure 3. A) WT PA1. B) WT PA2. C) *a2KO* PA1. D) *a2KO* PA2.

**Supplementary Figure S3.**
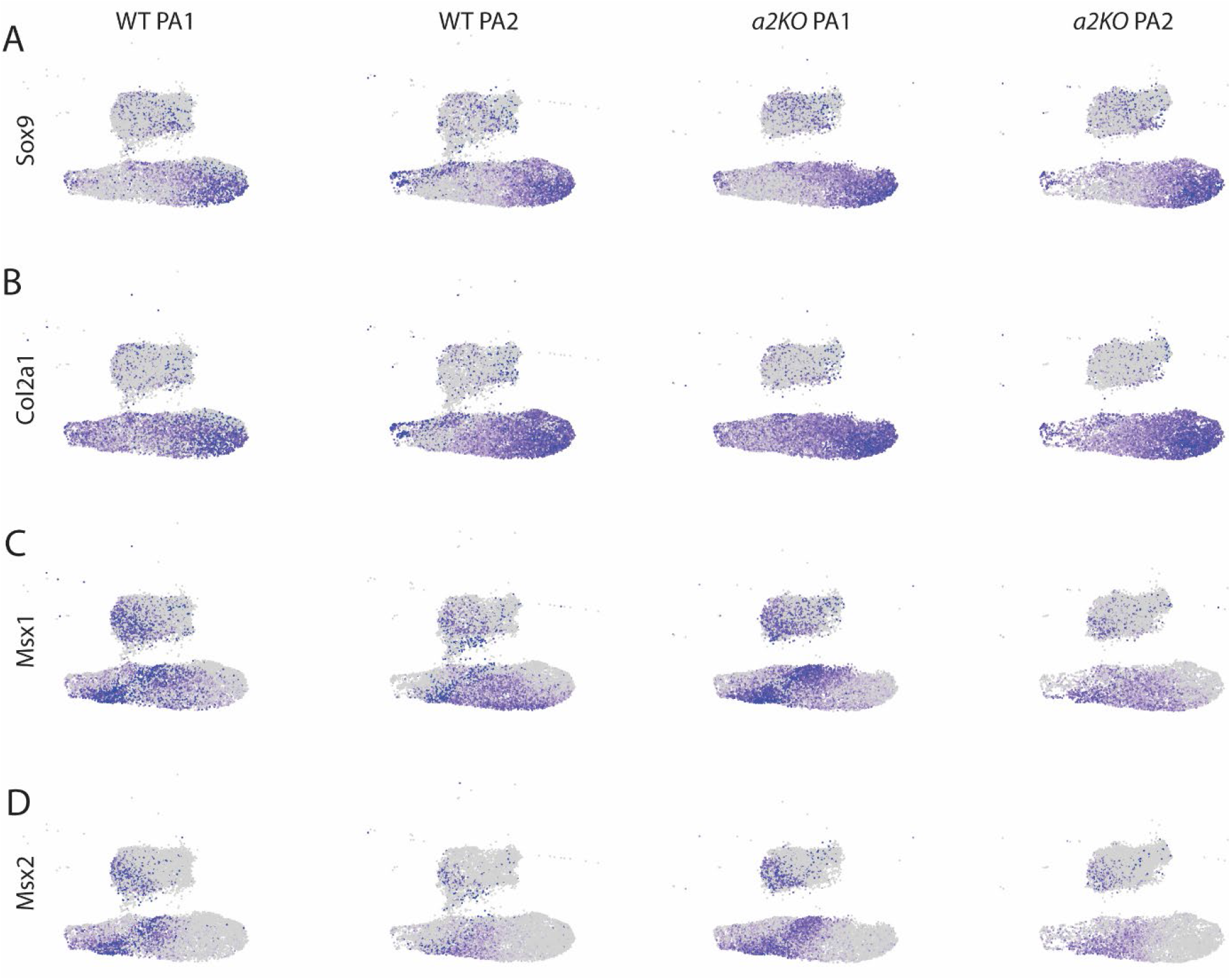
Visualization of chondrogenesis and osteogenesis markers across WT and *a2KO* PA1 and PA2. A) Chondrogenesis marker *Sox9*. B) Chondrogenesis marker *Col2a1*. C) Osteogenesis marker *Msx1*. D) Osteogenesis marker *Msx2*.

**Supplementary Figure S4.**
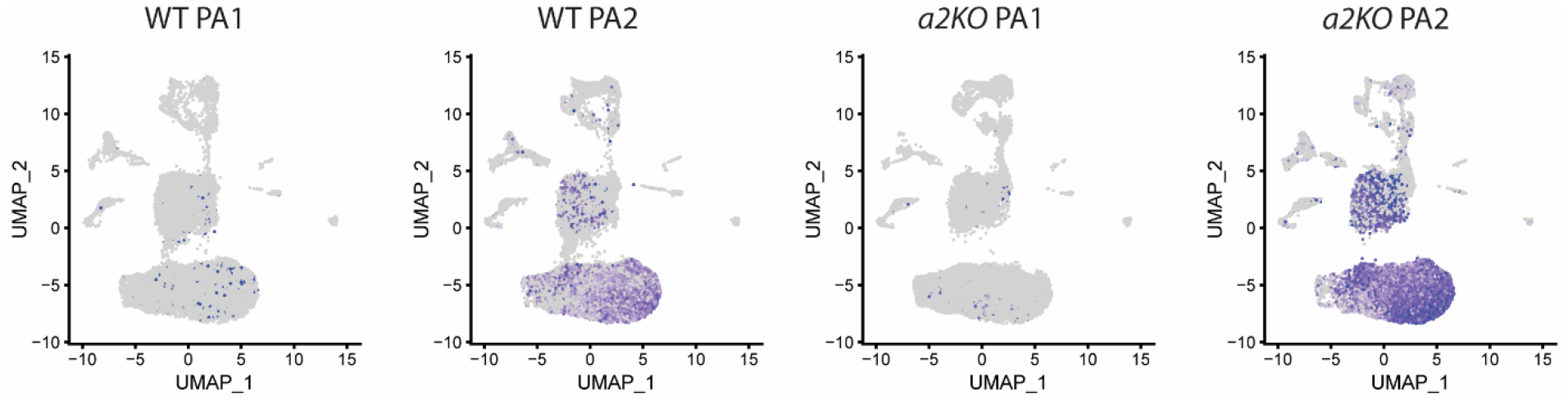
Visualization of *Hoxa2* expression in WT PA1, WT PA2, *a2KO* PA1, and *a2KO* PA2.

**Supplementary Figure S5.**
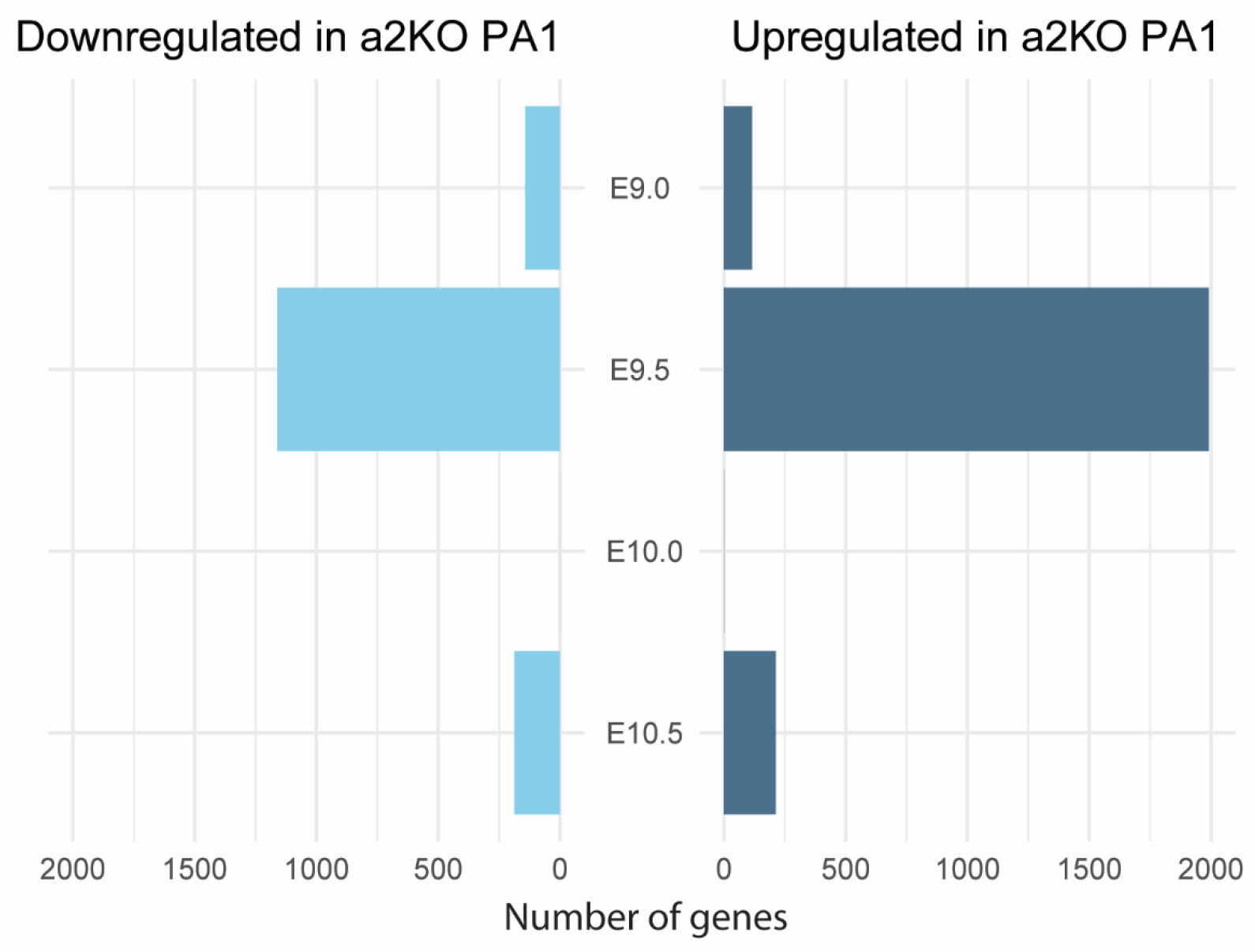
Bar plot showing genes differentially expressed in bRNAseq between WT PA1 and *a2KO* PA1 across timepoints.

